# *C9orf72*-associated G4C2 hexanucleotide repeat expression in *Drosophila* mushroom bodies causes age dependent TDP-43 pathology and dementia relevant phenotypes mediated in part by the glypican Dlp/GPC6

**DOI:** 10.64898/2026.05.08.723849

**Authors:** Brijesh S. Chauhan, Megan A. Brennan, Peter C. Forstmeier, Han Yifu, R. Keating Godfrey, Kendall Van Keuren-Jensen, Rita Sattler, Justin Ichida, Daniela C. Zarnescu

## Abstract

Hexanucleotide repeat expansions (HREs) in *C9orf72* are the most common genetic cause of amyotrophic lateral sclerosis (ALS) and frontotemporal dementia (FTD), yet the age-, sex-, repeat-length-, and circuit-specific influence on the pathology of neurons remains incompletely understood. Here, we established a *Drosophila* model of *C9orf72*-associated dementia by expressing G4C2 repeats in mushroom body neurons (MBNs), a brain region critical for memory, locomotion, and sleep. Expression of 44X G4C2 repeats ((G4C2)_44X_) led to progressive axonal thinning, age-dependent accumulation of Repeat Associated Non-AUG (RAN) translated GR-GFP dipeptide repeat (DPR) puncta, premature nuclear-to-cytoplasmic mislocalization of endogenous TDP-43, increased caspase, reduced lifespan and a loss of presynaptic active zones. Behaviorally, (G4C2)_44X_ expression caused locomotor hyperactivity, altered spatial working memory, and fragmentation of sleep architecture in an age- and sex-dependent manner, recapitulating core features of FTD. Surprisingly, the shorter (G4C2)_12X_ repeat, traditionally considered a control, also produced detectable RAN translation and intermediate phenotypes in aging MBNs, suggesting that length- and tissue-associated factors modulate repeat toxicity. We further identified a repeat-length- and age-dependent reduction of the glypican Dally-like protein (Dlp) in (G4C2)_44X_ consistent with disrupted Wnt-related signaling linked to TDP-43 proteinopathies. Restoring Dlp expression in MBNs mitigated locomotor and working-memory alterations, and loss of presynaptic active zones. In contrast, axonal degeneration, TDP-43 mislocalization, and lifespan were not significantly improved by restoring Dlp, suggesting that multiple mechanisms contribute to G4C2-induced toxicity. Supporting our findings in *Drosophila* MBNs, a CRISPRi screen in TDP-43 knock-down iNeurons identified GPC6, a human ortholog of Dlp, as a significant contributor to TDP-43 dependent synaptic loss. Together, our findings reveal an aging-sensitive, circuit-specific model of *C9orf72*-associated neurodegeneration and highlight roles for DPR accumulation and Dlp/GPC6 dependent synaptic loss in FTD pathomechanisms.

## Introduction

Mutations in the *C9orf72* gene that cause amyotrophic lateral sclerosis (ALS) and frontotemporal dementia (FTD) comprising expanded GGGGCC (G4C2) hexanucleotide repeats (HRs) are located within the first intron and represent the most common genetic abnormality in ALS (20%-50%) and FTD (10%-30%) [1, 2]. Expansions range from 24-30 to hundreds or thousands of G4C2 HRs in ALS/FTD brains, with heterogeneous distribution within and across tissues [3, 4]. In addition to typical ALS/FTD, HR expansions have been identified in Alzheimer’s disease, movement, and psychiatric disorders [5, 6]. Studies on the role of the G4C2 repeat expansions in ALS/FTD have indicated different disease-causing mechanisms for mutant *C9orf72*: *1)* toxic RNA gain-of-function; *2)* toxicity caused by Repeat-Associated Non-AUG initiated (RAN) translation, which leads to the accumulation of dipeptide repeat (DPR) proteins and *3)* haploinsufficiency [reviewed in 7, 8, 9]. RAN translation of G4C2 HR produces several DPRs [9, 10]; of these, arginine-rich DPRs (*i.e.*, GR, PR) are considered most toxic, as they disrupt RNA metabolism, ribosomal function, nucleo-cytoplasmic transport and proteostasis [11–14].

Clinically, the early stages of FTD are characterized by progressive cognitive and behavioral deterioration relative to baseline patient ability [9, 15]. Patients exhibit reduced short-term and working memory, emotional distress, and apathy [16] often coincident with increases in aggression, changes to sleep patterns, onset of wandering behaviors, and disinhibition of action and speech [17, 18]. With disease progression, many of the early symptoms worsen, and new symptoms may develop [18]. Despite these standard symptoms, behavioral changes in FTD are heterogenous and often vary with disease subtype, co-occurring motor neuron disease and sex [9, 15, 19, 20].

Previous *Drosophila* models of C9orf72 based on G4C2 HR expression in the retina, motor or all neurons have uncovered alterations in nucleo-cytoplasmic transport, mitochondrial function and translation [21–27], with one recent study showing that (G4C2)_44X_ HR expression in Kenyon cells (KCs) causes mislocalization of endogenous TDP-43 (TBPH) resembling human pathology [28]. Notably, single cell RNAseq (scRNAseq) of *Drosophila* brains expressing G4C2 HR panneuronally identifies KCs (referred herein as mushroom body neurons (MBNs)) as the most vulnerable neuronal population in the brain [29]. This circuit forms a layered network, reminiscent of vertebrate cortical neurons [30] and has been shown to be required for multi-modal and context-dependent learning [31, 32], including place learning [33]. The MBNs also regulate sleep [34, 35], social behavior [36], and gate behaviors attributed to other brain regions, such as decision-making [37] and aggression [38]. In addition to these FTD-relevant functions, existing *Drosophila* models of intellectual disability [39], Alzheimer’s disease (AD; [40]), and a recently reported model of dementia based on TDP-43 proteinopathy [41] have also targeted this circuit, highlighting the relevance of this circuit for modeling cognitive disorders.

Here, we leverage genetic and behavioral approaches to model *C9orf72*-associated dementia by expressing G4C2 HRs in MBNs. We found that a long repeat, (G4C2)_44x_ causes age-dependent axonal degeneration, DPR accumulation, TDP-43 nuclear to cytoplasmic mislocalization, increased caspase signaling, and a suite of dementia-relevant behavioral phenotypes, including locomotor, spatial working memory, and sleep deficits, as well as reduced lifespan and presynaptic active zones. Surprisingly, a shorter repeat ((G4C2)_12X_), originally developed as a control, and behaving as such in the retina [27], exhibited age-dependent RAN translation in MBs and caused similar phenotypes as the longer (G4C2)_44X_ repeats during aging. These findings suggest that RAN translation may be age and cell type specific and that DPR production mediates dementia-relevant *C9orf72* phenotypes. Overall, males exhibited more severe phenotypes compared to females, consistent with G4C2 sex-specific effects [42]. To gain insights into G4C2 HR’s mechanism of action, we investigated whether the glypican Dally-like protein (Dlp), a target of TDP-43 in flies and altered in ALS/FTD patient brains [41] could mediate aspects of *C9orf72*-associated phenotypes in the MB circuit. Indeed, Dlp was reduced in the context of (G4C2)_44X_ and restoring its expression in the MBNs rescued aspects of locomotor dysfunction and presynaptic active zones in an age dependent manner. Other phenotypes, including axonal thinning and TDP-43 mislocalization were not significantly mitigated, suggesting that Dlp mediates some but not all *C9orf72*-associated toxicity in the MB circuit. Bolstering the role of Dlp and its human ortholog GPC6 in synaptic formation and maintenance, we find that GPC6 knock-down worsens synapse loss in human iNeurons with TDP-43 loss of function, a molecular phenotype associated with *C9orf72* FTD. Taken together, these findings highlight several FTD relevant phenotypes caused by G4C2 HR expression in MBNs and identify Dlp/GPC6 as a modifier of *C9orf72*-associated phenotypes.

## Materials and Methods

### Drosophila genetics

*Drosophila melanogaster* stocks were cultured and maintained on standard cornmeal-agar medium at 25°C in 12-hour light/dark cycle. *C9orf72*-associated repeat expansion was modeled using previously described *w^1118^ P{w[+mC]=UAS-LDS-(G4C2)12.GR-GFP}2 ((G4C2)12X)* and *w^1118^;; P{w[+mC]=UAS-LDS-(G4C2)44.GR-GFP}9 ((G4C2)44X)* [27]. *w^1118^* was used as genetic background control. *w^1118^;P{w[+mC]=UAS-dlp.WT}3* was used to express *dally like protein* (*dlp*) [43]. For mushroom body neuron (MBN) specific expression, UAS lines were crossed to the split-Gal4 driver SS01276 [44]. *w^1118^; UAS-mCD8::RFP* served as a control reporter line. Recombinant lines co-expressing Dlp and mCD8::RFP were generated by standard genetic techniques between *w^1118^; P{w[+mC]=UAS-dlp.WT}3* and *y[1] w[*]; P{w[+mC]=UAS-mCD8.mRFP.LG}10b* and validated by immunofluorescence. Similarly, recombinant lines co-expressing (G4C2)_44X_ and Dlp were generated by recombining *w^1118^; P{w[+mC]=UAS LDS-(G4C2)44.GR-GFP}9* with *w^1118^;P{w[+mC]=UAS-dlp.WT}3*. A complete list of *Drosophila* lines and corresponding RRIDs is provided in Table S1.

### Immunofluorescence

*<u>Drosophila brains</u>* were dissected in cold HL3 saline (70 mM NaCl, 5 mM KCl, 22 mM MgCl_2_, 10 mM NaHCO_3_, 5 mM trehalose, 15 mM sucrose, and 5 mM HEPES, pH 7.3) followed by fixation in 4% paraformaldehyde (PFA, Santa Cruz Biotechnology sc-281692) for 60 minutes at room temperature (RT). Brains were washed three times with 1X PBS, pH 7.4 (CORNING 46-013-CM) at RT for 10 min each. Next, the brains were permeabilized in PBS (Sigma-Aldrich T9284) with 0.1% Triton™X-100 (0.1% PBS-T) for 30 minutes at RT, followed by additional permeabilization with 0.3% PBS-T for 20 minutes at RT on slow speed orbital shaker. Following treatment with blocking solution (5% normal goat serum (Abcam ab7481), 2% Bovine serum albumin (Sigma-Aldrich A9647-506) in 0.3% PBS-T) for 45 minutes at RT, the brains were incubated with primary antibodies for 48 hours at 4°C. Mouse anti-Fasciclin-II (1:40, DSHB 1D4) was used to detect mushroom body axons; rabbit anti-TBPH (1:100), a kind gift from Fen-Biao Gao, UMass Chan Medical School) was used to detect endogenous TDP-43 (dTDP-43) [45]; and mouse anti-Dlp (1:5, DSHB 13G8) was used to detect Dally like protein, mouse anti-BRP (1:100, DSHB nc82) stained active zones [46]. After incubation with primary antibodies, the brains were washed three-times with 0.3% PBS-T at RT for 15 minutes, then treated with secondary antibodies: goat anti-mouse IgG Alexa 633 (1:500, Invitrogen A-11005), or goat anti-rabbit IgG Alexa 594 (1:500, Invitrogen A-11012) and goat anti-mouse IgG Alexa 594 (1:500, Invitrogen A-11005) or goat anti-mouse IgG Alexa 488 (1:500, Invitrogen A-11001). GFP was detected with goat anti-GFP-FITC (1:300, Rockland 800-656-7625) incubated overnight at 4°C. To detect DNA, brains were incubated with Hoechst (1:10,000 in PBS, Invitrogen H3570), for 10 minutes at RT following treatment with secondary antibodies. Brains were mounted in 80% glycerol and 2% propyl-gallate. Images of MBNs were acquired using a Leica TCS SP8 confocal inverted microscope S/N 8100001634 with an HC PL APO 40X/1.30 oil CS2 objective. Brains were imaged using Z series with slices captured every 1-2 µm.

*<u>iPSC-derived neurons</u>* were fixed in 4% PFA for 20 min at 4°C, followed by permeabilization with 0.1% Triton X-100 in PBS for 10 min at room temperature 8 days after siRNA transfection. Cells were blocked in PBS containing 10% donkey serum and 1% bovine serum albumin (BSA) for 1 hour at room temperature and subsequently incubated with primary antibodies overnight at 4°C. Cells were then washed with 0.1% PBS-T three times and incubated with Alexa Fluor-conjugated secondary antibodies (Life Technologies) for 1 hour at room temperature. Cells were then washed with 0.1% PBS-T three times and mounted on slides with Vectashield (Vector Labs). Images were acquired on an LSM 800 confocal microscope with oil immersion at 63x (Zeiss). The following primary antibodies were used: chicken anti-MAP2 (1:2000, Abcam, ab5392) and mouse anti-Synapsin (1:1,000, Synaptic Systems 106 011).

### Image analyses

All image analyses were performed in FIJI v.2.17.0/Image J unless otherwise noted.

#### Mushroom body morphology

At least 6 brains/age/genotype/sex were used for analyses throughout lifespan (1-3, 30, and 60 days). FasII maximum intensity projections were used to measure axonal lobe thickness throughout the lifespan. Width was measured in three locations along the lobe and averaged. This was performed for both sides of the circuit and averaged per brain. The average axonal thickness for G4C2 HR-expressing brains was normalized to the w^1118^ genetic background controls. Statistical significance was determined using Kruskal-Wallis with Dunn’s multiple comparisons tests following removal of outliers with the ROUT test in GraphPad Prism 10.6.1.

#### Measurement of GR-GFP puncta numbers and sizes

GR-GFP puncta number and size in the γ lobes were quantified as described [41]. At least 6 brains /age/sex/genotype were analyzed. Slices from each Z-stack image of MBs were used, selected from the top, middle, and end of each stack. The γ lobes were traced and the area measured. DPR (GR-GFP) puncta were quantified using FIJI’s particle analysis tool within the traced area. The values from the three individual slices were averaged for each γ lobe and across both lobes. Statistical significance was determined using Kruskal-Wallis with a Dunn’s multiple comparisons test in GraphPad Prism 10.6.1.

#### Nuclear versus total dTDP-43 (TBPH) intensity

Subcellular localization of endogenous dTDP-43 (TBPH) in MBNs was measured from the middle of a Z stack comprising all channels (Hoechst, GFP, and TBPH). MBN nuclear regions of interest (ROIs) were manually defined based on Hoechst then whole-cell body ROIs, identified by the presence of GFP, were drawn using the freehand selection tool. Mean TBPH fluorescence intensity was measured within each ROI in the appropriate channel. Due to MBNs having very thin cytoplasms, to improve accuracy, TBPH signal was measured in whole cells, including nuclei [41]. TBPH nuclear to whole-cell ratios were calculated for each cell and used as a measure of TBPH nuclear to cytoplasmic mislocalization. Approximately 6-8 MBNs were analyzed per brain, with at least six brains per sex and genotype included in analyses.

#### DCP1 puncta density and size quantification

A central slice was selected from each MBN Z stack, and the area of the MBNs was traced. Cell number, area and mean DCP1 signal were measured. The DCP1 mean signal was normalized to the number of MBNs.

#### Dlp expression in MB γ lobes

The γ lobe was outlined using GFP fluorescence or Dlp expression if no GFP signal was present (*e.g.*, w^1118^ and 12X controls). The contrast was adjusted to clearly visualize the γ lobe and mean Dlp intensity was measured. Background Dlp signal was subtracted using a nearby area to yield corrected Dlp fluorescence values. Both γ lobes were analyzed and averaged for each brain (N**∼**6 brains/sex/genotype). Statistical analyses were performed using one-way ANOVA, with Dunnett’s multiple comparisons test in GraphPad Prism 10.6.1.

#### Active zone quantification

The MB γ lobe was manually outlined as a region of interest (ROI), and the mean fluorescence intensity of nc82 signal was measured from three representative optical sections (top, middle, and bottom of Z stack), which were averaged to obtain the final value for each brain. Background intensity was measured from a nearby region outside of the MBs and subtracted from the ROI signal. Background corrected nc82 signal intensity was calculated as: γ lobe mean intensity - background mean intensity, and analyzed using the Mann-Whitney unpaired test to determine statistical significance in GraphPad Prism 10.6.1.

#### Synaptic density in iPSC neurons

Synapsin puncta were counted by particle analysis and normalized to 100 μm neurite length [detailed in 47] traced using the NeuronJ plugin. Neurite analysis was performed by manually counting branching neurites from single neurons with MAP2 staining.

### Behavioral studies

#### Spatial learning and locomotion behavioral assays

The spatial learning and memory function of adult flies was assessed using Y-maze assays, as previously described [41]. Flies aged 1-3, 30 or 60 days were tested during peak activity (17:00-19:00) in miniature Y-mazes (N∼50/sex/genotype/age). Following a two-minute acclimation period, alternation and movement behavior was evaluated over 10 one-minute intervals using the EthoVision XT software (v11.5). Each one-minute bin was considered a technical replicate. Statistical differences were evaluated using Kruskal-Wallis with Dunn’s multiple-comparison test, and values more than ±2 standard deviations from the median were considered outliers and excluded from subsequent analyses. Five parameters were measured in the one-min bins, including the (1) total distance moved, (2) the alternation score, which is the number of actual alternations over the maximum potential alterations, (3) the number of actual alternations over the total distance moved, (4) the velocity of the flies when they are in motion, and (5) the frequency of breaks in movement.

#### Sleep analysis

Sleep and activity were assessed using the *Drosophila* Activity Monitor (DAM) System 3 (Trikinetics). Adult flies (N∼32/sex/genotype/age) from three age time points (1-3, 30, and 60 days) were monitored using the Activity Monitor (DAM) for 7 days at 1-minute intervals. Individual-aged flies were placed in DAM tube containing fresh food in a 25 °C incubator with a 12-hour Light-Dark (LD) cycle. The count of beam breaks was collected per minute with sleep bouts defined as a period of > 5 minutes with no beam breaks. Data were collected over the first three full days, and the days were considered technical replicates. Data from biological replicate trials were pooled. Except for the actograms, all activity and sleep bouts and counts were separated by LD status and log1p normalized before analysis. Statistical differences were evaluated using Kruskal-Wallis with Dunn’s multiple-comparison test, and values lying more than ± 2.0 standard deviations from the median were considered outliers and excluded from subsequent analyses.

### Lifespan analysis

Virgin flies of both sexes were collected and monitored for deaths daily over their lifespan. N equal or > 100 flies were used per genotype and sex, pooled from two independent replicates. Food was changed weekly, and flies were stored at a standard 12-hour light/dark cycle in a 25 °C incubator. Lifespan data was collected in Excel and analyzed in GraphPad Prism 10.6.1. using a Kaplan-Meier survival test with the Gehan-Breslow-Wilcoxon test to evaluate differences between lifespan curves. The same data set was evaluated for differences in mean lifespan using Kruskal-Wallis with Dunn’s multiple-comparison test.

### iPSC-Derived Neurons

Human induced pluripotent stem cells (iPSCs) constitutively expressing a doxycycline-inducible *NGN2* cassette were used for neuronal differentiation, as previously described [48]. iPSCs were dissociated and resuspended in mTeSR (STEMCELL 85850) medium consisting of 10 μM Y-27632 (Sellechchem S1049). Cells were seeded onto Matrigel (Corning 354230) coated plates at 50 x 10^3^ per well (6-well format, 2 mL per well) on day 0. Medium was switched to N2 medium consisting of DMEM/F12 (Gibco 10565042) supplemented with 1% MEM non-essential amino acids (Gibco 11140076), 1% N2 supplement (Gibco 17502048), 1% Pen-Strep (Corning 30001CI), 10 ng/mL NT-3 (Peprotech 450-03), 10 ng/mL BDNF (R&Dsystems 11166-BD), and 1 μg/mL doxycycline (Cayman 14422) to induce *NGN2* expression for 2 days. On day 3, pre-differentiated cells were dissociated with accutase (STEMCELL 07920) and replated in neuronal maturation medium consisting of Neurobasal (Gibco 21103049) supplemented with 1% MEM non-essential amino acids, 1% GlutaMAX (Gibco 35050079), 1% Pen-Strep, 2% B27 supplement (Gibco 17504044), 10 ng/mL NT-3, 10 ng/mL BDNF, 1 μg/mL laminin (Gibco 17504044), 10 μM Y-27632 and 1 μg/mL doxycycline. Cells were plated onto plates coated with Poly-ʟ-Lysine (Sigma P6282) and laminin at a density of 50 x 10^3^ cells per cell (24-well format, 500 uL per well). Maturation medium was half-changed every other day on day 5 without Y-27632. Doxycycline was removed from Maturation medium on day 9. On day 7, primary cortical mouse glia was dissociated with accutase and plated onto neuronal culture in Maturation medium at a 1:5 astrocytes to seeded neurons ratio [49].

### siRNA transfection

siRNAs were added with Lipofectamine RNAiMAX Transfection Reagent (Invitrogen 13778075) on day 19 induced neurons in Maturation medium [detailed in 49]. Induced neurons were treated with a scrambled siRNA (Invitrogen 4390846) or a TARDBP siRNA (Invitrogen 4392420) at 10 nM. A complete medium change was done on the following day. Cell cultures were harvested for qRT-PCR 48 hours later using primers for GAPDH (CGAGATCCCTCCAAAATCAA; GTCTTCTGGGTGGCAGTGAT) and TARDBP (TCATCCCCAAG CCATTCAGG; TGCTTAGGTTCGGCATTGGA).

### Molecular cloning and viral production

sgRNAs for GPC6, GAPDH, EEF2, ACTB, RPL10L and non-targeting sgRNAs [as described in 50] were cloned into a lentiviral plasmid (Addgene 84832) for gRNA expression with Genscript. Viruses were produced as follows. HEK293T cells were transfected in a 10-cm dish at 80-90% confluence with viral vectors containing each sgRNA and viral packaging plasmids (pPAX2 and VSVG for lentivirus) using polyethylenimine (PEI) (Sigma-Aldrich). The medium was changed 24 hours after transfection. Viruses were harvested at 48 hours and 72 hours after transfection. Viral supernatants were filtered with 0.45-μm filters, incubated with Lenti-X concentrator (Clontech) for 24 hours at 4°C, and centrifuged at 1,500 g at 4°C for 45 minutes. Pellets were resuspended in DMEM plus 10% FBS (200 μL per 10-cm dish of HEK293T) and stored at –80°C.

sgRNAs used for each targeting gene and non-targeting controls are listed below:

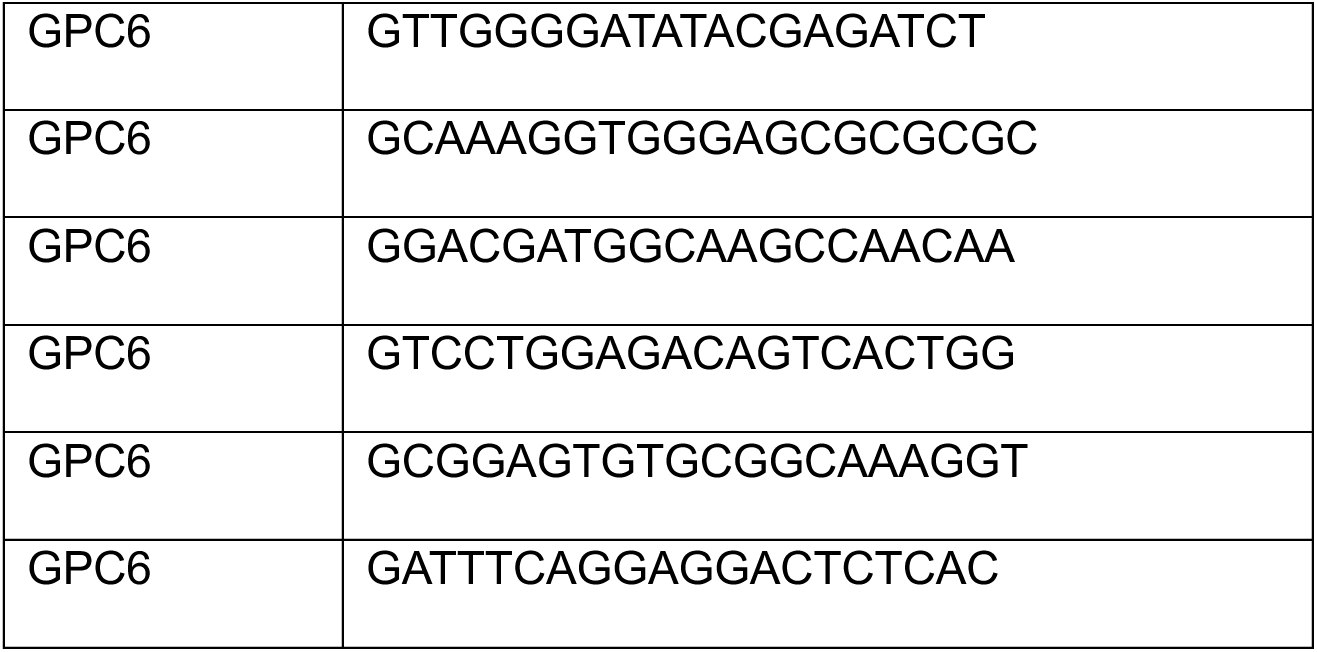
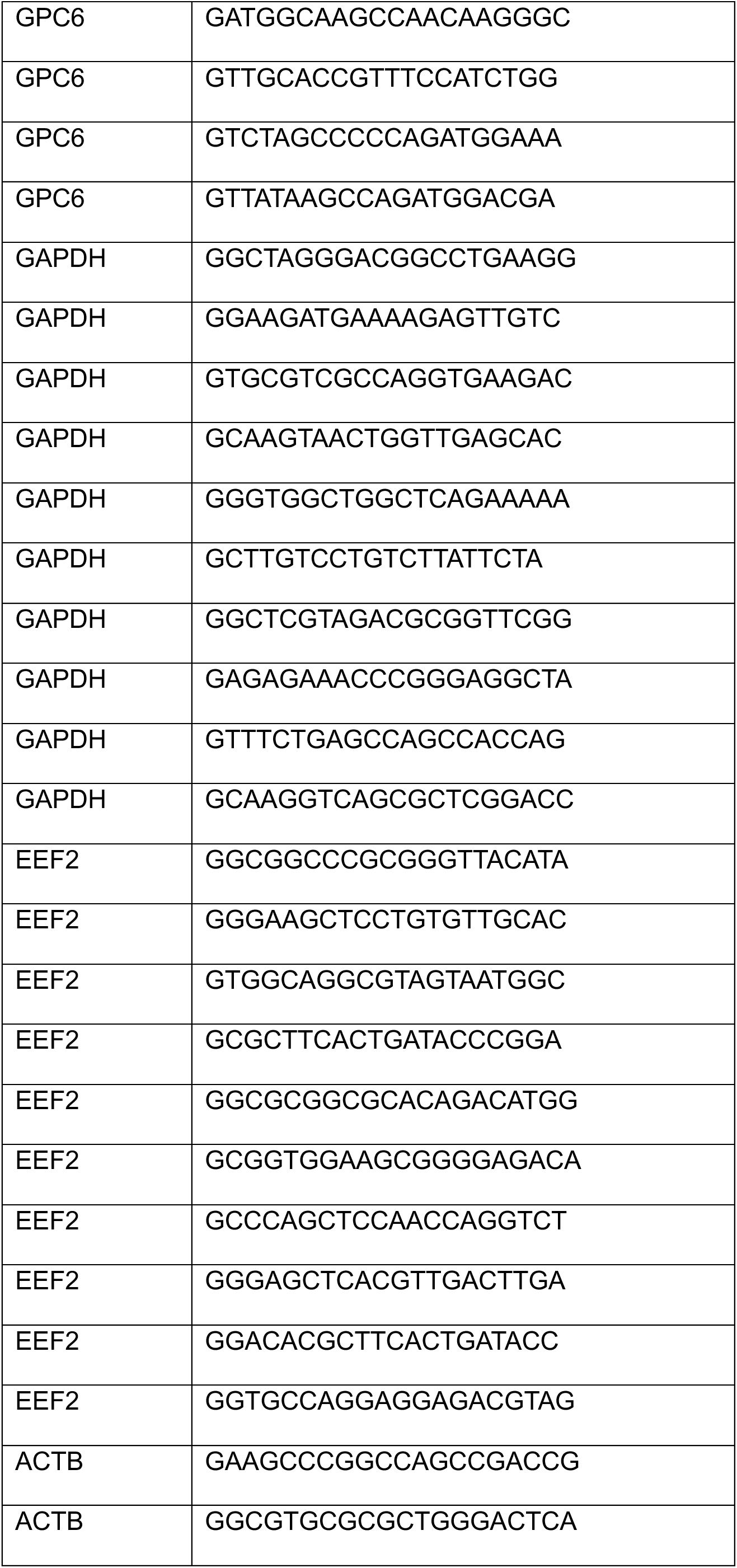
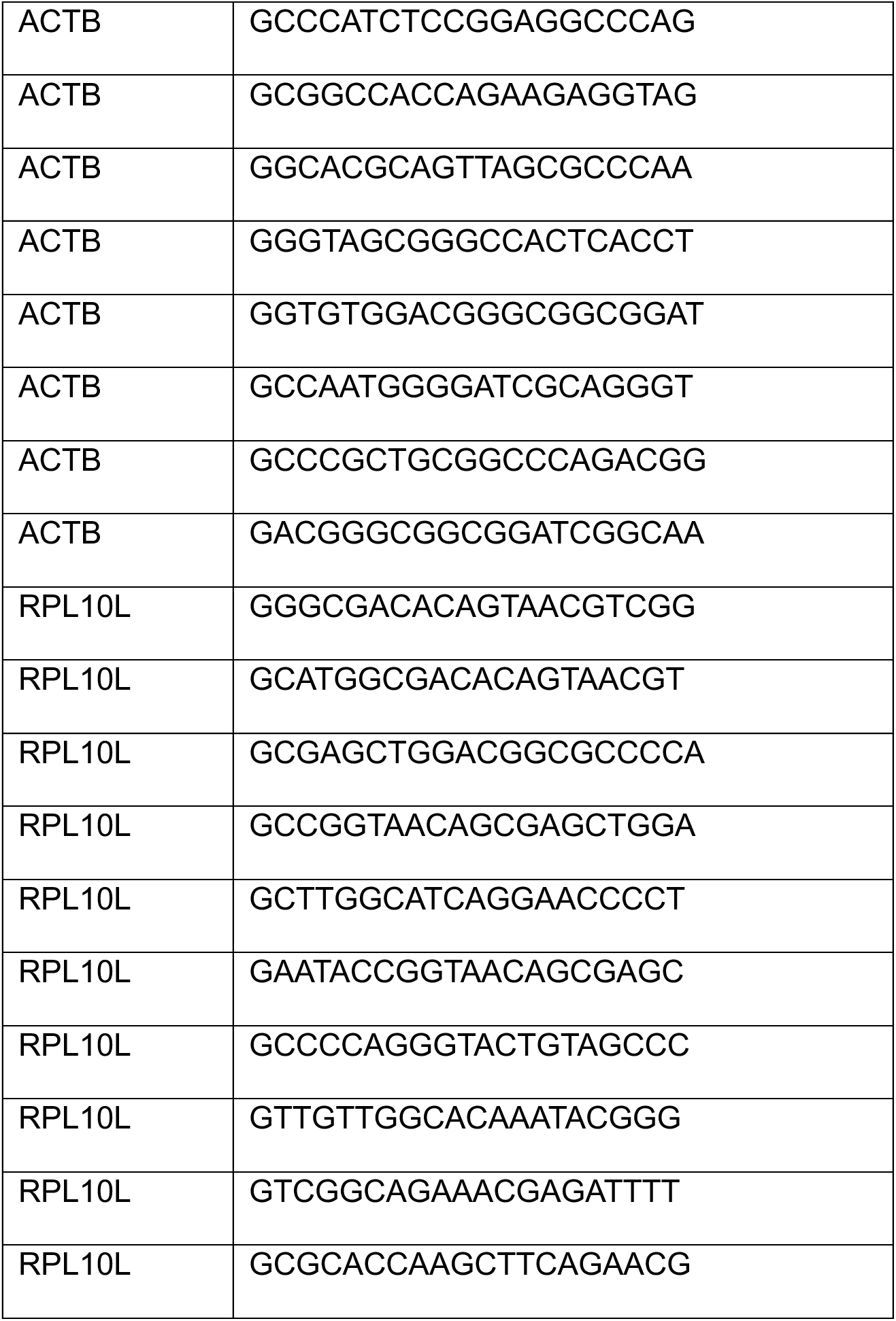

### Rabies virus trans-neuronal tracing

For trans-neuronal tracing, pre-differentiated cells were plated onto 96-well plate (200 μL per well) in Maturation medium at a density of 20 x 10^3^ cells per well. On day 4 of iPSC-derived neuron culture, neurons were transduced in Maturation medium with AAV virus to express TVA, mCherry and G protein (Addgene 104330). A complete medium change was done on the following day. AAV was titered to observe 10% of mCherry positive cells against DAPI in cultured neurons 7 days after AAV transduction. On day 21 of iPSC-derived neuron culture, neurons were transduced in Maturation medium with rabies virus (Addgene 32635) to express EnvA-G-deleted rabies and GFP. Rabies virus was obtained from Salk Viral Vector Core. A complete medium change was done on the following day. Longitudinal tracking of mCherry and GFP expressing neurons was performed using Molecular Devices ImageXpress before Rabies transduction and every other day after transduction for 15 days. For each rabies tracking assay, iPSC-derived neurons were quantified from three technical independent differentiations treated with either a scrambled siRNA or a TARDBP siRNA on day 19 of neuronal differentiation as described above. Trans-neuronal quantification was performed by counting mCherry negative and GFP positive neurons.

### sgRNA enrichment assay

For GPC6 knocking down evaluation, all sgRNAs described above were equally mixed in lentiviral suspension and added to iPSCs seeded on a 6-well plate in mTESR medium consisting of 8 μg/mL polybrene and 10uM Y-27632 at 100 x 10^3^ per well. A complete media change with mTESR medium consisting 1μg/mL puromycin was performed on next day. Puromycin treatment proceeded until 80% of iPSC express BFP compared to un-transduced control cells assessed by flow cytometry. iPSCs were then seeded for rabies virus trans-neuronal tracing assay as described above. On day 16 after rabies transduction, neurons were incubated in Papain (Worthington LK003176) with DNase1 (Worthington LK003170) reconstituted per manufacturer instructions for 30 minutes at 37⁰C and dissociated by gentle pipetting as described in [51]. Single-cell suspension was pelleted and resuspended in ice-cold Neurobasal supplied with 0.2% bovine serum albumin (BSA) and filtered through 40 μM Flowmi tips (Sigma BAH136800040). Single cells were then sorted with BD ARIA sorter to collect GFP positive and mCherry negative cells. Sorting was gated based on control neurons without rabies transduction. Genomic DNA of sorted cells were purified using Monarch® Genomic DNA Purification Kit. sgRNA amplicon was enriched by PCR reaction with 5’ Truseq primer and 3’ common primer (CAAGCAGAA GACGGCATACGAGATCGACTCGGTGCCACTTTTTC). Sample indices for demultiplexing were added in the 5’ Truseq primers (TARDBP siRNA: aatgatacggcgaccaccgagatctacacgatcggaagagcacacgtctgaactccagtcacCTTGTAgcacaaaaggaaactcaccct; scramble siRNA: aatgatacggcgaccaccgagatctacacgatcggaagagcacacgtctgaactccagtcacGCCAATgcacaaaaggaaactcaccct). gRNA amplicon samples were pooled and sequenced with on an Illumina NextSeq 500 as described [52]. Raw count of each sgRNA was analyzed using ScreenProcessing [53]. sgRNA enrichment analysis was performed using the Robust Rank Aggregation (RRA) algorithm with MAGeCKFlute [54]. sgRNA-level log2 fold changes and depletion P-values were calculated using MAGeCK test. MAGeCK models sgRNA read counts using a negative binomial–based framework after library-size normalization. *p.value* represents the P-value for depletion or enrichment of an sgRNA in the treatment condition compared with control. Gene-level positive and negative selection were assessed using MAGeCK robust rank aggregation/α-RRA. sgRNAs were first ranked by P-values estimated from MAGeCK’s negative-binomial count model, and gene-level significance was calculated by testing whether sgRNAs targeting each gene were non-randomly enriched among the most depleted or enriched sgRNAs.

### Statistical analyses

Statistical analyses were performed using GraphPad Prism 10.6.1. unless otherwise noted, and described for individual methods.

## Results

### G4C2 hexanucleotide repeat (HR) expression in mushroom body neurons (MBNs) causes age-dependent axonal thinning in adult brains

To study the effects of G4C2 HR expression on the MB circuit we used a previously generated (G4C2)_44X_ repeat transgene that includes a leader sequence (LDS) of 114-base pairs upstream of the repeat in intron 1 of *C9orf72* within ALS/FTD patient genomes, and a GFP cloned in frame with GR dipeptides generated via RAN translation [27]. This HR line reports GR translation via the GFP tag and affords a more patient-relevant context, as the LDS is thought to influence pathological mechanisms. A shorter (G4C2)_12X_ HR and *w^1118^* were used as controls for the longer 44X repeat and for the genetic background, respectively (see Materials and Methods for details). To evaluate the effect of *C9orf72*-associated HRs in the MB circuit, the G4C2 transgenes were expressed in MBNs using a split-GAL4 driver that restricts expression to MBNs forming the α/β and γ lobes [SS012761, 41, 44]. We examined the morphology of MBNs using immunostaining for FasII, an axonal membrane marker [55]. These experiments reveal a visible thinning of MB axons with age in the context of G4C2 HR expression and indicate cell-type specific progression of thinning. The γ lobes already show significant thinning in young flies (**Figure 1A-I2**, **Figure K**: P < 0.0001 for 1-3 DO, P < 0.0001 for 30 days old (DO), P = 0.0002 for 60 DO), whereas the α lobes exhibit comparable morphology with controls in young and middle aged flies (1-3 and 30 DO, respectively) (**Figure 1A-I2**: **1J** P = 0.9999 for 1-3 DO, P = 0.0520 for 30 DO) but exhibit a significant reduction in axonal thickness at the oldest time point (**Figure 1J**: P <0.0001 for 60 DO). Similar results were observed for β lobes, axons belonging to the same cells as the α lobes (Supplemental **Figure S1)**. The expression of (G4C2)_12X_ repeats had no significant effects on axonal thickness throughout the MB circuit (**Figure 1A-K**). Our experiments showed overall similar G4C2-associated phenotypes in male and female flies, albeit males often showed somewhat more severe phenotypes, suggesting sex dependent effects. For simplicity, we have chosen to show the male data in the main text figures and the female data in the Supplement. Overall, these results indicate that the expression of (G4C2)_44X_ expanded repeats in the MB circuit causes age-dependent axonal thinning. Interestingly, γ neurons, which are born before the α and β neurons, exhibit more severe age-dependent phenotypes, highlighting the aging component of *C9orf72* ALS/FTD.

**Figure 1.**
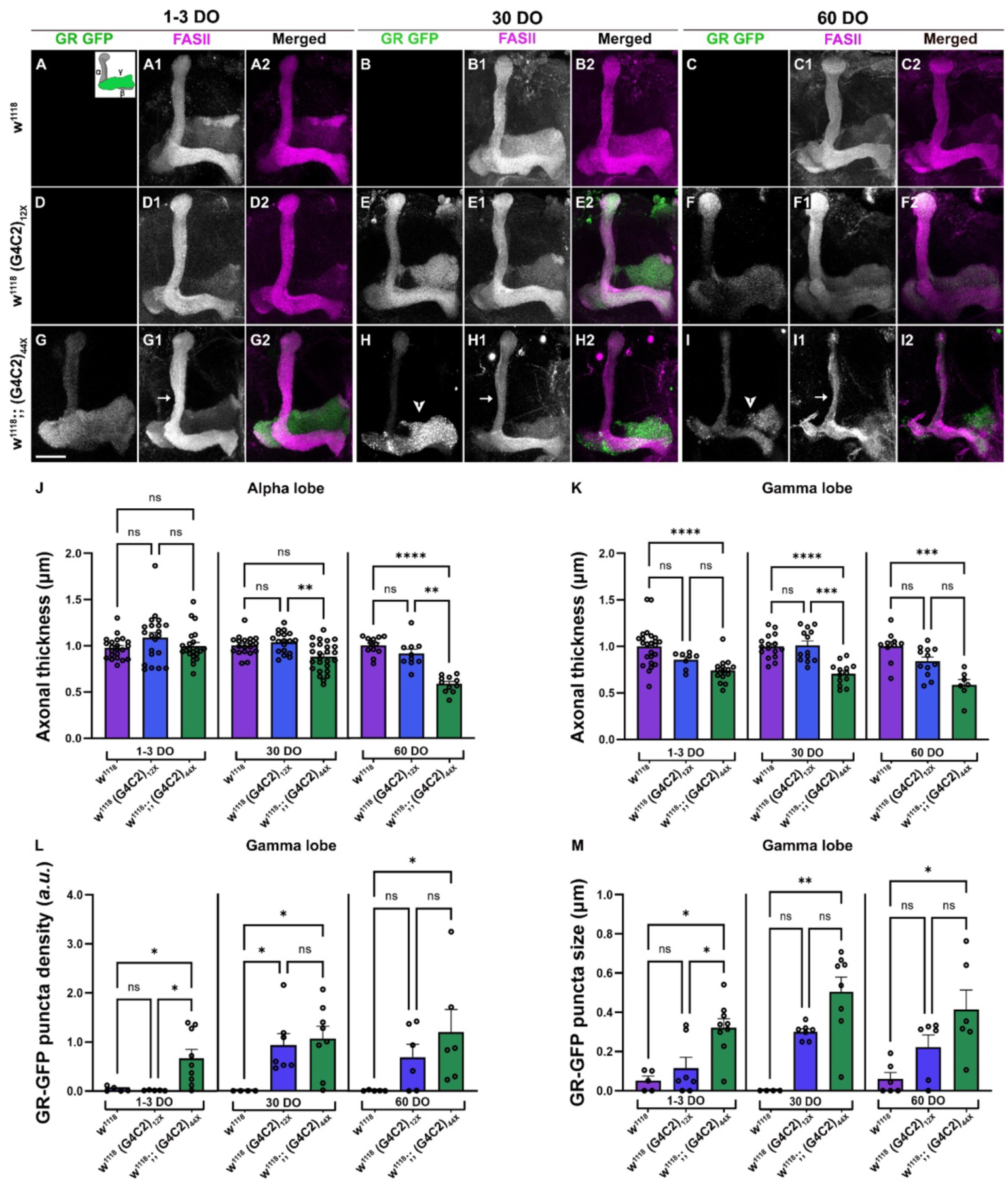
(G4C2)_44X_ expression causes age dependent axonal thinning in the MB circuit. **(A-C2)** w^1118^ controls stained for GR-GFP and FasII. α, β, γ lobe cartoon shown in **(A). (D-F2)** (G4C2)_12X_ stained for GR-GFP and FasII. **(G-I2)** (G4C2)_44X_ stained for GR-GFP and FasII. GR-GFP **(A-I)**, FasII **(A1-I1)** and merged images **(A2-I2)**, as indicated. Arrows indicate thinning α lobes in (G4C2)_44X_, and arrowheads indicate GR-GFP puncta in γ lobes. **(J, K)** Quantification of axonal thickness in 1-3 DO, 30 DO, and 60 DO adult MBs. Data were normalized to w^1118^ controls. **(L, M)** GR-GFP puncta density in γ lobes **(L)**. GR-GFP puncta size in γ lobes **(M)**. Genotypes and ages, as indicated. Scale bar: 50 μm. Kruskal-Wallis with Dunn’s multiple comparison test was used to determine statistical significance; ^ns^P>0.05, *P<0.05, **P<0.01, ***P<0.001, ****P<0.0001.

### RAN translation and GR GFP puncta accumulation are age-dependent in the mushroom body circuit

While expressing G4C2 HRs in MBNs, we were able to detect robust GFP signal as a reporter of RAN translation in (G4C2)_44X_ flies (**Figure 1A-I2**). Interestingly, while (G4C2)_12X_ short repeat behaved as a control when measuring axonal thinning (**Figure 1A-K**) and showed no GR-GFP signal in young flies (**Figure 1D-D2**, 1-3DO), GR-GFP expression was detected in 30 and 60 DO adults, suggesting that RAN translation may be enhanced during aging (**Figure 1E-F2**). To determine when RAN translation products are first detected, we imaged the GFP tagged GR products signal before 30 DO and found expression as early as 5 DO (data not shown). Next, we quantified the puncta density and size of GR-GFP particles and found that there are significantly more and larger puncta in the context of (G4C2)_44X_ HR expression compared to w^1118^ genetic background controls (**Figure 1L**: P = 0.0237 for 1-3 DO, P = 0.0107 for 30 DO, P = 0.0196 for 60 DO; **Figure 1M**: P = 0.0319 for 1-3 DO, P = 0.0014 for 30 DO, P = 0.0154 for 60 DO). In contrast, (G4C2)_12X_ HRs showing intermediate levels of GR-GFP puncta compared to w^1118^ (**Figure 1L**: P > 0.9999 for 1-3 DO, P = 0.0330 for 30 DO, P = 0.1257 for 60 DO; **Figure 1M**: P > 0.9999 for 1-3 DO, P = 0.1756 for 30 DO, P = 0.2760 for 60 DO). Given that previously published data did not show (G4C2)_12X_ expression in the retina of young flies [27], our results show for the first time the production of RAN translation products from the shorter 12X repeat that may be specific to the MB circuit and/or age-dependent. Since the (G4C2)_12X_ HR flies are similar to controls in some phenotypic assays and resemble (G4C2)_44X_ HR-like phenotypes in other assays, presumably due to the expression of RAN translation products, we are highlighting (G4C2)_44X_ associated phenotypes in comparison with w^1118^ controls in the main text, and report the findings from all genotypes, including the (G4C2)_12X_ short repeat, in the supplement.

### Endogenous *Drosophila* TDP-43 (TBPH) exhibits premature nuclear to cytoplasmic mislocalization in the context of G4C2 HR expression in MBNs

A hallmark of pathology across a large proportion of *C9orf72* ALS/FTD is the presence of cytoplasmic TDP-43 aggregates accompanied by loss of nuclear TDP-43 [56]. To determine whether G4C2 HR expression causes TDP-43 pathology, we examined the expression of endogenous TDP-43 (TBPH) in MBNs. These experiments show that young flies (1-3 DO) expressing cytoplasmic GFP as a control exhibit mostly nuclear TBPH while (G4C2)_44X_ expression causes significant mislocalization of TBPH to the cytoplasm (**Figure 2A-A3, D-D3** and **G**: P < 0.0001). (G4C2)_12X_ expression exhibits an intermediate level of TBPH mislocalization (see Supplemental **Figure S2** for all genotypes and female data). Interestingly, although at 30 DO, controls expressing cytoplasmic GFP still exhibit mostly nuclear TBPH, by 60 DO, TBPH is localized increasingly to the cytoplasm, suggesting that the localization of endogenous TDP-43 itself is regulated during lifespan. Quantification of TBPH localization indicates that (G4C2)_44X_ HR expression causes mislocalization compared to controls (**Figure 2B-B3**, **E-E3**, **G**: P < 0.0001 for 30 DO, **Figure C-C3**, **F-F3**, **G**: P =0.0036 for 60 DO). Taken together, these observations in 44X flies are consistent with premature mislocalization of endogenous TBPH, highlighting a potentially accelerated aging process in disease.

**Figure 2.**
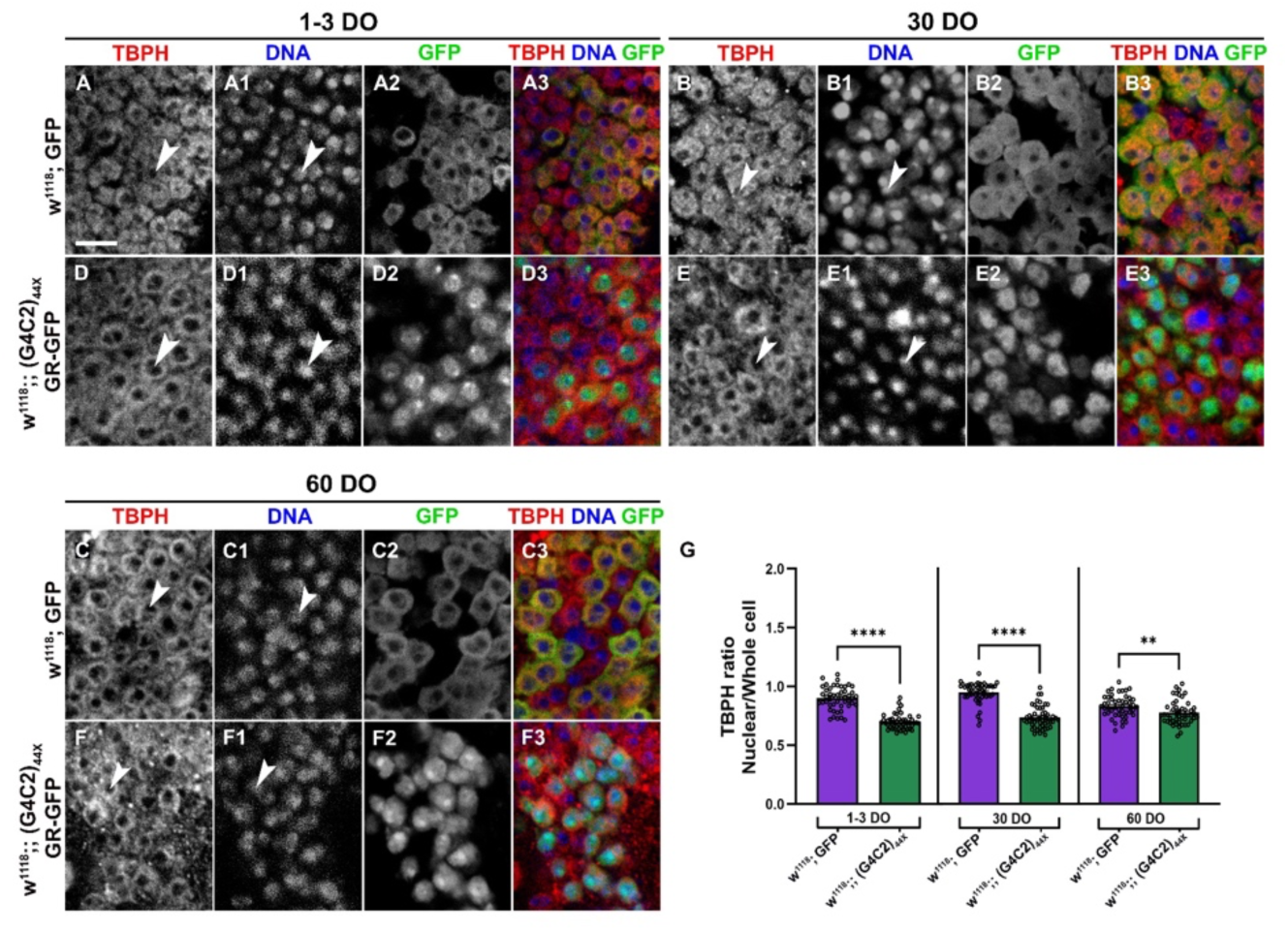
G4C2 HR expression in MBNs alters endogenous TDP-43 (TBPH) localization in an age-dependent manner. Representative confocal images of *Drosophila* MBNs at 1-3 DO **(A-A3, D-D3)**, 30 DO **(B-B3, E-E3)**, and 60 DO **(C-C3, F-F3)**. Each age group shows TBPH immunostaining **(A-F)**, nuclear DNA labeled with Hoechst **(A1-F1)**, GFP fluorescence **(A2-F2)**, and merged images **(A3-F3)**. Scale bar: 5 µm. **(G)** Quantification of TBPH localization in KCs across lifespan. Kruskal-Wallis with Dunn’s multiple-comparison test; *P < 0.05, **P < 0.01, ****P < 0.0001.

### (G4C2) HR expression in MBNs causes an age-dependent increase in caspase signaling

Next, we tested whether (G4C2) HR expression caused cell death signaling, which is associated with TDP-43 pathology and neurodegeneration [57]. To this end, we examined the expression of cleaved caspase 1 (DCP1), a core component of the apoptotic signaling pathway, in MBNs, in the context of G4C2 HR expression compared with controls expressing cytoplasmic GFP throughout the lifespan. These experiments showed that while a small number of cells express DCP1 in controls at old age (60 DO), (G4C2)_44X_ expression exhibits a progressively higher DCP1 signal when normalized to cell number (**Figure 3A-F2**: P = 0.4841 for 1-3DO, P = 0.0152 for 30 DO, P = 0.0264 for 60 DO), while (G4C2)_12X_ exhibits intermediate levels of DCP1 (Supplemental **Figure S3**). These findings suggest that G4C2 HRs cause an increase in caspase signaling in a repeat length- and age-dependent manner.

**Figure 3.**
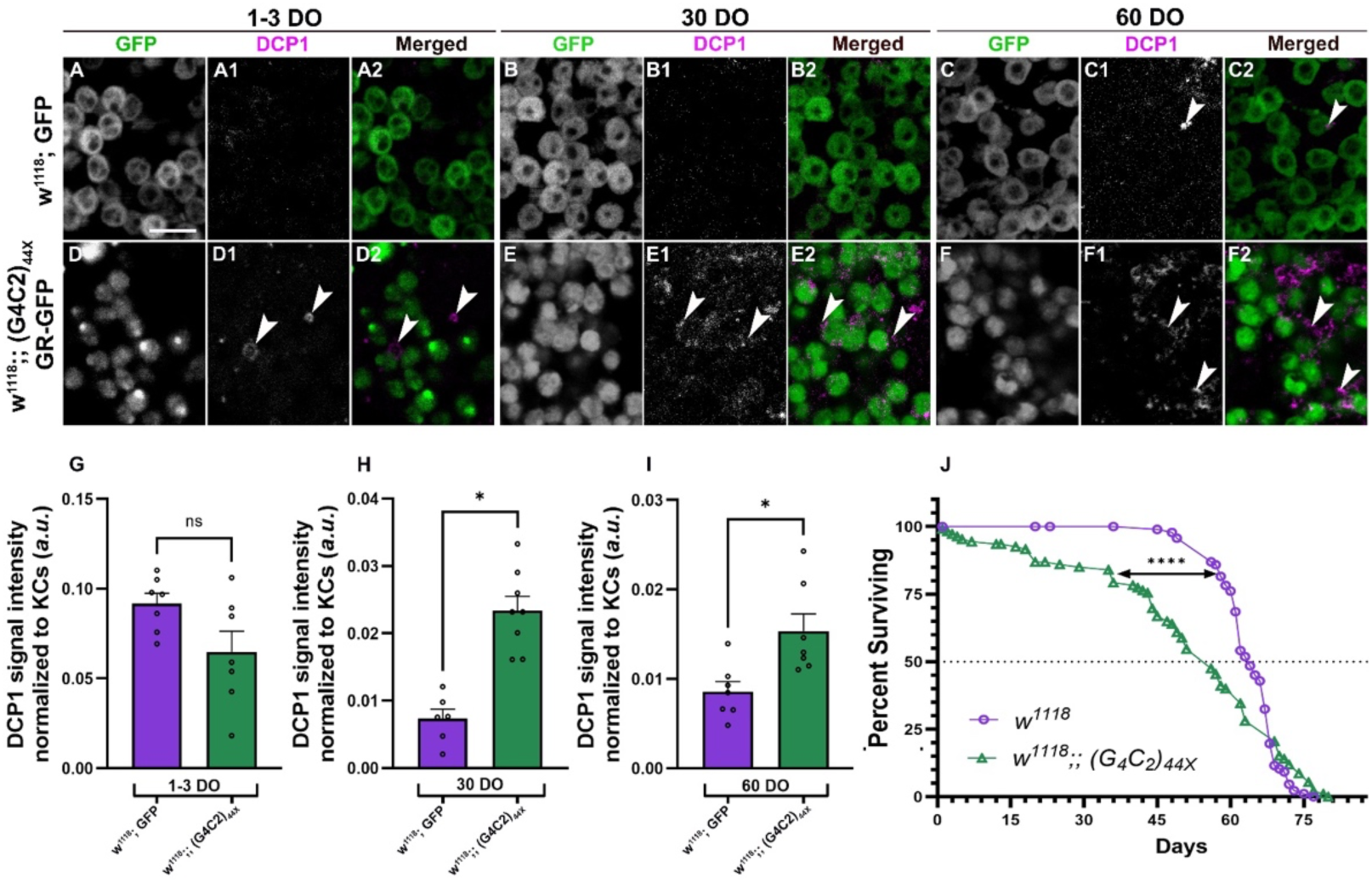
G4C2 HR expression in *Drosophila* MBNs induces age-dependent increase in caspase signaling in MBNs and reduces lifespan. (A – F2) Confocal images showing GFP expression. **(A - F)** and cleaved DCP-1 immunostaining **(A1 – F1)** in KCs of (G4C2)_44X_ and GFP expressing male flies. Genotypes, as indicated on the left. Ages, as indicated on the top: 1-3 DO, 30 DO, and 60 DO. Merged images, as shown **(A2 - F2)**. Scale bar: 5 µm. **(G - I)** Quantification of cleaved DCP-1 signal intensity in 1-3 DO **(G)**, 30 DO **(H)**, and 60 DO **(I)** brains. Data were analyzed using Kruskal-Wallis with Dunn’s multiple-comparison test; * = P < 0.05. **(J)** Survival curves of male expressing G4C2 repeat expansions in MBNs. The dotted line indicates the 50% survival point (median lifespan). N ≥ 90 flies /genotype/sex. Data were analyzed using the Kaplan-Meier method. Statistical significance was assessed by the Gehan-Breslow-Wilcoxon test; ****P < 0.0001.

### G4C2 HR expression in MBNs reduces lifespan

We previously found that human TDP-43 overexpression in MBNs causes a small, but significant decrease in lifespan [41]. To determine whether C9-associated G4C2 HRs may have a similar effect, we measured the lifespan of G4C2-expressing adults compared to the w^1118^ genetic background controls. These experiments showed that both 44X and 12X repeats of G4C2 cause a small but significant reduction in lifespan (**Figure 3J** and Supplemental **Figure S3**, respectively). (G4C2)_44X_ HR expressing male flies showed reduced average lifespan (50.29 days +/- 1.950) as compared with w^1118^ (63.64 days +/- 0.641; P = 0.0200) (**Figure 3J**). Females also experienced a shorter average lifespan from 77.98 +/- 0.6826 days for the w^1118^ to 69.17 +/- 1.549 days for (G4C2)_44X_ (P =0.0002) (Supplemental **Figure S3**). In this assay, (G4C2)_12X_ males and females exhibited a similar reduction in average lifespan as (G4C2)_44X_ when compared to w^1118^ controls (49.8 days +/- 1.118, P <0.0001, N ≥90 male flies and 68.87 +/- 1.038 days, P <0.0001) (Supplemental **Figure S3**). Taken together, these findings show that the expression of G4C2 HR in MBNs causes a small but statistically significant decrease in average lifespan, similar to the effects of FTD in human patients [58] and consistent with the MB circuit playing a role in longevity [59].

### G4C2 HR expression in MBNs causes changes in locomotor behavior and spatial working memory defects in an age-, sex-, and time-dependent manner

We recently reported a model of FTD based on TDP-43 proteinopathy that showed alterations in spatial working memory [41]. To test whether G4C2 HR expression causes similar alterations in working memory and locomotor behaviors, we used miniature Y-mazes, adapted from rodent studies, to measure spontaneous alternation behavior, a measure of working memory [60], by scoring the number of three consecutive arm entries over a set period (see Materials and Methods and **Figure 4A**). These experiments showed that male (G4C2)_44X_ expressing flies exhibit increased locomotor activity throughout lifespan (**Figure 4B**: P < 0.0001 for 1-3DO, P < 0.0001 for 30 DO, P = 0.020 for 60 DO) consistent with some patient symptoms [17, 61] and degradation of the MBs [62]. Significantly higher alternation scores were observed in 1-3 DO and 30 DO flies (**Figure 4C**: P = 0.0009 for 1-3 DO, P = 0.0496 for 30 DO), suggesting that (G4C2)_44X_ expression causes alterations in working memory. Interestingly, 60 DO (G4C2)_44X_ expressing male flies showed similar alternation scores as controls. This may be due to aging effects on the MB circuit, which could obscure our ability to detect certain behavioral deficits. To measure the potential impact of locomotor behavior on alternations, we normalized the number of spontaneous alternations to movement and found a similar trend throughout lifespan (compare **Figures 4C and D**). Next, to more closely characterize the increased movement behavior of G4C2 HR expressing flies, we analyzed moving velocity, which was higher in (G4C2)_44X_ compared to w^1118^ controls in the 1-3 DO and 30 DO male flies (**Figure 4E**: P < 0.0001 for 1-3 DO, P < 0.0001 for 30 DO, and P > 0.9999 for 60 DO). This increase in speed was accompanied by a decrease in movement breaks throughout lifespan (**Figure 4F**: P < 0.0001 for 1-3 DO, P < 0.0001 for 30 DO and for 60 DO), suggesting that (G4C2)_44X_ expressing male flies exhibit longer bursts of rapid movement compared to controls. This locomotor phenotype resembles FTD-related hyperactivity observed in patients [63].

**Figure 4.**
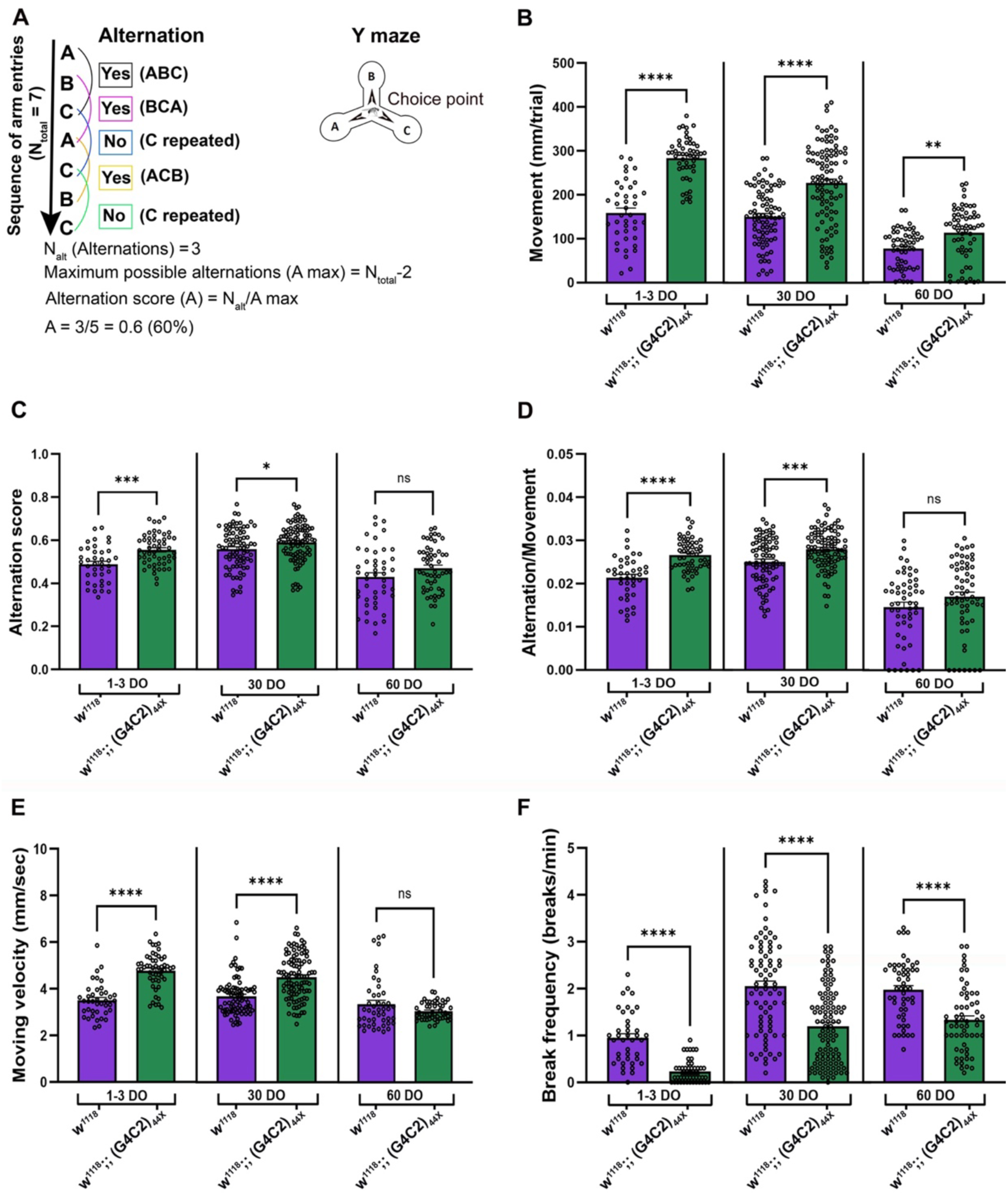
Spatial working memory of (G4C2)_44X_ expressing male *Drosophila* at different age points. **(A)** Schematic representation of the Y-maze and spontaneous alternation measurements. Alternation scores were calculated as the number of alternations (Nalt) divided by the number of maximum possible alternations. **(B)** Movement. **(C)** Alternation scores indicating spontaneous alternations. **(D)** Number of Alternations normalized to movement. **(E)** Moving velocity, and **(F)** Break frequency. Genotypes and ages, as indicated. N≈35 flies/sex/genotype/timepoint. Significance was calculated using Kruskal-Wallis with Dunn’s multiple-comparison test; *P < 0.05, **P < 0.01, ***P < 0.001, ****P < 0.0001.

Interestingly, (G4C2)_44X_ expressing female flies showed less robust phenotypes than the males, suggesting that (G4C2)_44X_ expression confers sex-specific effects (Supplemental **Figure S4**). Meanwhile, both male and female (G4C2)_12X_ expressing flies exhibited changes that are variable throughout lifespan, including female and male locomotor hypoactivity (Supplemental **Figure S4**). Taken together, these findings indicate that G4C2 HR expression, particularly longer (G4C2)_44X_, causes increased locomotion and changes in spatial working memory that are consistent with FTD-like phenotypes, and these changes occur in an age- and sex-specific manner.

### Expression of the (G4C2)_44X_ HR in the *Drosophila* MBNs alters daytime and nighttime sleep

Given reports of FTD-associated sleep behavioral deficits [64, 65], we assessed our flies for changes in sleep and activity. Using the well-established *Drosophila* Activity Monitor (DAM) assays [66], we monitored flies for several days and found that HR expression caused changes in sleep and activity patterns throughout the lifespan of the flies. Young male flies show altered sleep both during the day and night, with a reduction in bout numbers (**Figure 5A**: 1-3 DO daytime P < 0.0001; **Figure 5C**: 1-3 DO nighttime, P < 0.0001) and an increase in bout length (**Figure 5B**: 1-3 DO daytime, P < 0.0001; **Figure 5D**: 1-3 DO nighttime, P < 0.0001). In contrast, 30 and 60 DO male flies generally exhibited an increase in bout numbers (**Figure 5A**: 30 DO daytime, P < 0.0001, 60 DO daytime, P < 0.0001; **Figure 5C**: 30 DO nighttime, P =0.0099, 60 DO nighttime, P =0.0102) and a decrease in bout length (**Figure 5B**: 30 DO daytime, P < 0.0001, 60 DO daytime, P <0.0001; **Figure 5D**: 30 DO nighttime, P =0.0950, 60 DO nighttime, P = 0.0071). Qualitatively consistent with these findings, overall activity across zeitgeber time appears to be altered in (G4C2)_44X_ flies during lifespan, with 1-3 DO males appear to show reduced activity (**Figure 5E**), while 30 and 60 DO males appear to show an increased amount of activity as measured by the number of DAM beam breaks (**Figures 5F**, **G**). Similar results were obtained when measuring sleep using DAMs in female flies expressing (G4C2)_44X_ HRs compared to controls, while (G4C2)_12X_ expressing flies showed variable behaviors across the lifespan (Supplemental **Figure S5**). Taken together, these results indicate that the expression of disease-causing (G4C2)_44X_ HRs can produce both sleep consolidation in young flies and sleep fragmentation during aging, similar to reported sleep disturbances in FTD patients [65], indicating age dependent G4C2 HR-associated disease-like phenotype.

**Figure 5.**
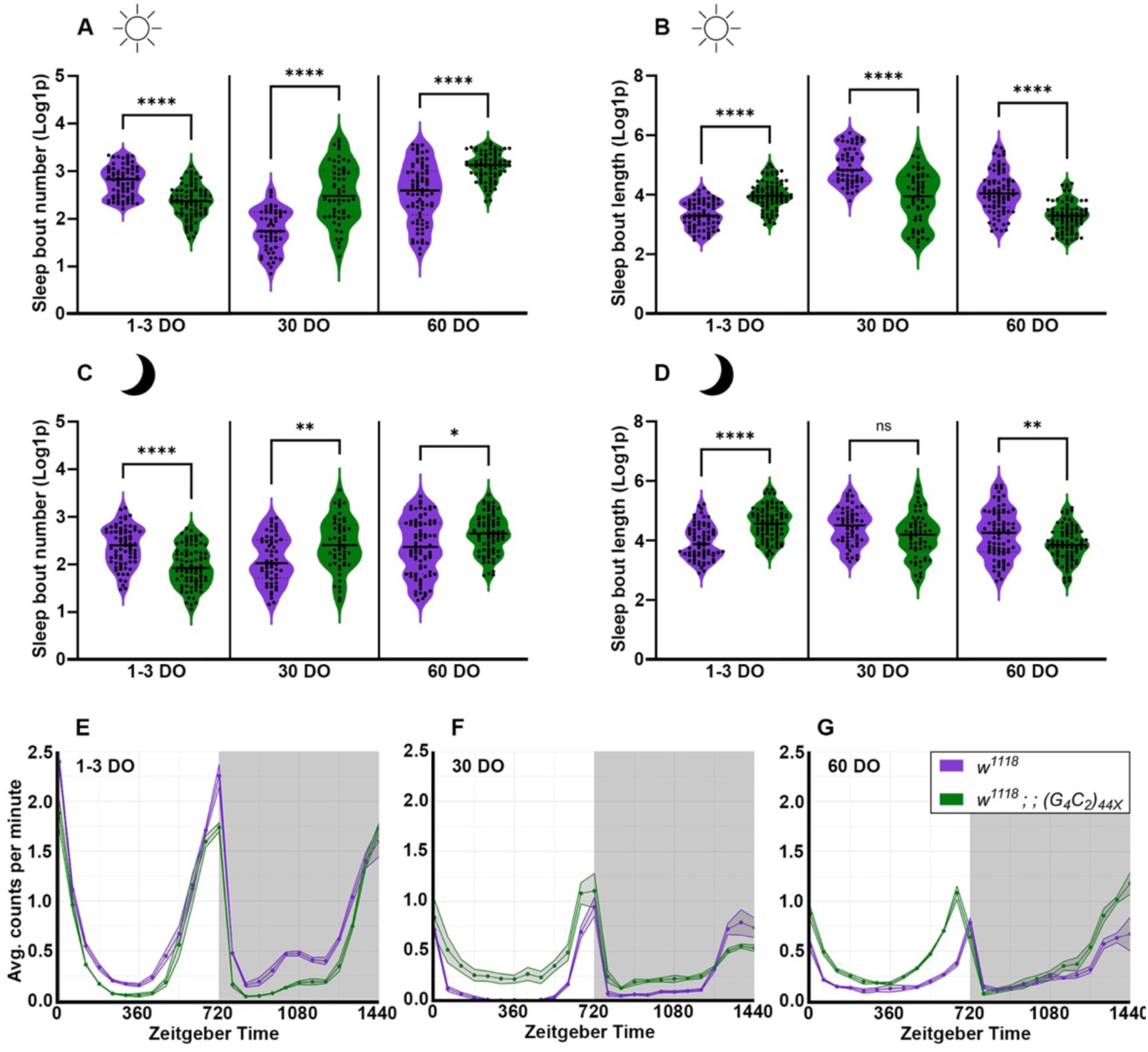
Sleep behavior and activity patterns are altered by the expression of (G4C2)_44_X HR. **(A, B)** Sleep bout numbers **(A)** and sleep bout lengths **(B)** during the 12-hour light period (daytime)**. (C, D)** Sleep bout numbers **(C)** and sleep bout lengths **(D)** during the 12-hour dark (nighttime) cycle. **(E-G)** The average number of beam breaks across zeitgeber time for 1-3 DO **(E)**, 30 DO **(F)**, and 60 DO **(G)** male flies. Nighttime indicated by a gray background. N∼32/genotype/age. Significance was determined using a Kruskal-Wallis ANOVA with Dunn’s test for multiple comparisons; *P < 0.05, **P < 0.01, ***P < 0.001, ****P < 0.0001.

### G4C2 HR expression in MBNs causes age-dependent reduction of the glypican Dally-like protein (Dlp)

We previously reported that *dlp* mRNA is enriched in TDP-43 complexes and TDP-43 proteinopathy in MBNs reduces Dlp expression with ageing [41]. Dlp and its human orthologs GPC6/GPC4 are glypicans that regulate wingless (Wg/Wnt) signaling with recent reports showing that Dlp is altered in both *Drosophila* models and patient tissues with TDP-43 pathology [67] and dysregulated GPC6 contributes to nuclear pore and TDP-43 dysfunction in mouse models [68]. Furthermore, snRNA seq experiments showed that *GPC6* mRNA is altered in neurons with TDP-43 pathology within *C9orf72* FTD brains [69]. Given the alterations in endogenous TDP-43 expression we found in the context of our *C9orf72* model (**Figure 2**), we sought to determine whether similar changes in Dlp occur in MBNs expressing G4C2 HRs as reported in models of TDP-43 proteinopathy. Immunofluorescence experiments showed that while in the γ lobe axons of young male flies (1-3 DO), Dlp expression was comparable between (G4C2)_44X_ and controls (see arrowheads in **Figures 6A-A2**, **D-D2**, **G**: P = 0.5559), it was significantly reduced in (G4C2)_44X_ at 30 DO and 60 DO relative to controls, demonstrating genotype and age dependent decline (see arrowheads in **Figure 6 B-B2**, **E-E2**, **G**: P = 0.0033 for 30 DO, and **Figure 6C-C2**, **F-F2**, **G**: P = 0.0008 for 60 DO). Male flies expressing short repeats (G4C2)_12X_ showed no significant changes at any age (Supplemental **Figure S6**). No changes in Dlp were observed in MBN cell bodies, suggesting that the effect is primarily localized to the axonal compartment (Supplemental **Figure S6**). Similar to males at 1-3 DO, female flies expressing either short or long G4C2 HRs exhibited no significant effect on Dlp. At 30 DO, 12X repeat females were still comparable to controls while 44X showed a significant reduction in Dlp expression. By day 60, Dlp levels were significantly reduced by long HRs in females. These results are consistent with repeat size- and age-associated vulnerability in females (Supplemental **Figure S6**) and demonstrate a repeat length and age-dependent reduction of Dlp expression in MB γ lobe axons, with longer repeats accelerating the decline.

**Figure 6.**
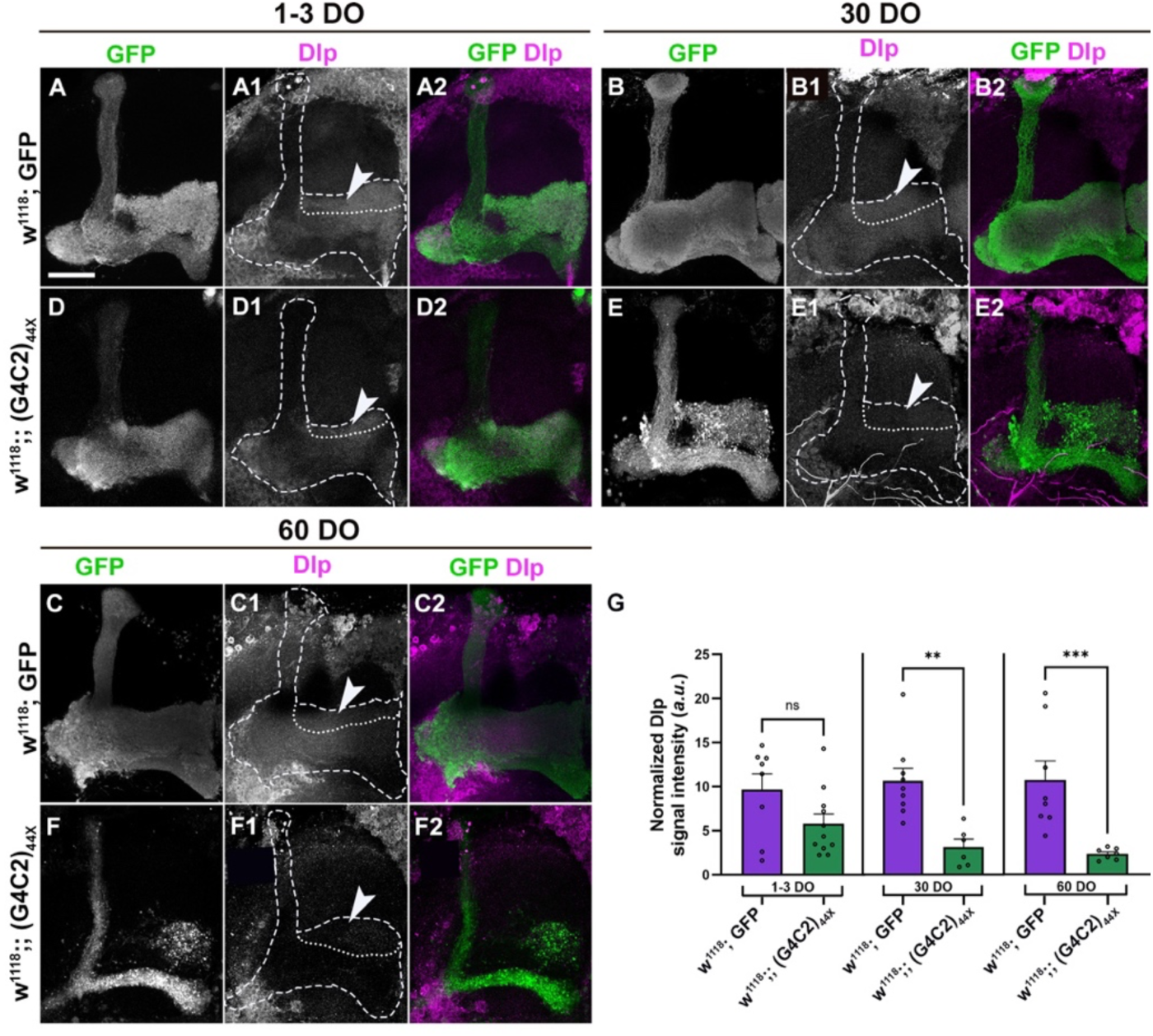
Age-dependent reduction of Dally-like protein (Dlp) expression in MB γ lobes expressing (G4C2)_44X_ HRs. **(A-F2)** GFP and Dlp expression in the MB axons at 1-3 DO **(A-A2, D-D2)**, 30 DO **(B-B2, E-E2)**, and 60 DO **(C-C2, F-F2)**. Genotypes, as indicated on the left. Stainings and ages, as indicated on top. Arrowheads indicate γ lobes where Dlp was quantified. **(G)** Quantification of Dlp fluorescence intensity in γ lobes across ages. N=6-11 male flies/genotype/age. Statistical significance was determined using Kruskal-Wallis with Dunnett’s multiple comparisons test; **P < 0.01, ***P < 0.001Scale bar: 50 µm.

### Restoring Dally-like protein (Dlp) expression in MBNs mitigates behavioral deficits in (G4C2)_44X_ Drosophila

The observation of age- and (G4C2)_44X_ dependent reduction in Dlp levels within γ lobes (**Figure 6**) and our previous findings showing that restoring Dlp expression can mitigate phenotypes associated with TDP-43 proteinopathy in motor neurons and MBNs [41, 67] led us to hypothesize that overexpressing Dlp may also rescue (G4C2)_44X_ associated phenotypes. To test this, we conducted miniature Y-maze assays to measure locomotion and spontaneous alternation in flies co-overexpressing Dlp and (G4C2)_44X_ (Dlp (G4C2)_44X_ co-OE) compared to (G4C2)_44X_ or Dlp overexpression (OE) alone in MBNs. These experiments confirmed that male (G4C2)_44X_ expressing flies exhibit increased locomotor activity in 1-3 DO and 30 DO, consistent with previous results (**Figure 4B**). While Dlp and (G4C2)_44X_ co-overexpression (co-OE) has no significant effect on locomotor activity in 1-3 DO flies (**Figure 7A**: P <0.0001 for Dlp OE vs (G4C2)_44X_, P < 0.0001 for Dlp OE vs Dlp (G4C2)_44X_ co-OE, P = 0.5518 for (G4C2)_44X_ vs Dlp (G4C2)_44X_ co-OE), it partially rescues this phenotype in 30 DO flies (**Figure 7A**: P < 0.0001 for Dlp OE vs (G4C2)_44X_, P = 0.0057 for Dlp OE vs Dlp (G4C2)_44X_ co-OE, P = 0.0068 for (G4C2)_44X_ vs Dlp (G4C2)_44X_ co-OE). (G4C2)_44X_ associated alternation score increases were mitigated in both 1-3 DO (**Figure 7B**: P = 0.0051 for Dlp OE vs (G4C2)_44X,_ P = 0.9999 for Dlp OE vs Dlp (G4C2)_44X_ co-OE, P = 0.0271 for (G4C2)_44X_ vs Dlp (G4C2)_44X_ co-OE) and 30 DO (**Figure 7B**: P = 0.78994 for Dlp OE vs (G4C2)_44X,_ P = 0.0090 for Dlp OE vs Dlp (G4C2)_44X_ co-OE, P < 0.0001 for (G4C2)_44X_ vs Dlp (G4C2)_44X_ co-OE). It is interesting that Dlp OE alone resembles (G4C2)_44X_ alternation score phenotype at 30 DO, suggesting that precise levels of Dlp are critical for proper physiology [67]. Closer characterization of locomotor behaviors showed that (G4C2)_44X_ expressing flies recapitulated increased moving velocity at 1-3 DO and 30 DO (**Figure 4E**), however Dlp co-OE had no impact on this phenotype at either age tested (**Figure 7C**: P < 0.0001 for Dlp OE vs (G4C2)_44X_ 1-3 DO, P < 0.0001 for Dlp OE vs Dlp (G4C2)_44X_ co-OE 1-3 DO, P = 0.8232 for (G4C2)_44X_ vs Dlp (G4C2)_44X_ co-OE 1-3 DO, P = 0.0222 for Dlp OE vs (G4C2)_44X_ 30 DO, P = 0.0033 for Dlp OE vs Dlp (G4C2)_44X_ co-OE 30 DO, P > 0.9999 for (G4C2)_44X_ vs Dlp (G4C2)_44X_ co-OE 30 DO). Meanwhile, (G4C2)_44X_ expression-associated decrease in movement breaks throughout lifespan was recapitulated (**Figure 4F**) and rescued by Dlp co-OE at 1-3 DO and 30 DO (**Figure 7D**: P < 0.0001 for Dlp OE vs (G4C2)_44X_ 1-3 DO, P = 0.1551 for Dlp OE vs Dlp (G4C2)_44X_ co-OE 1-3 DO, P = 0.0007 for (G4C2)_44X_ vs Dlp (G4C2)_44X_ co-OE 1-3 DO, P < 0.0001 for Dlp OE vs (G4C2)_44X_ 30 DO, P = 0.0959 for Dlp OE vs rescue 30 DO, P = 0.0171 for (G4C2)_44X_ vs Dlp (G4C2)_44X_ co-OE 30 DO). To allay concerns of potential GAL4-UAS dilution effects caused by the presence of two UAS-driven transgenes in Dlp (G4C2)_44X_ co-OE compared to one UAS when (G4C2)_44X_ is expressed alone, we used western blotting to evaluate GR-GFP expression and found it to be comparable between flies expressing (G4C2)_44X_ with or without Dlp co-OE (Supplemental **Figure S7**). In addition to Y-maze assays, we also probed the ability of Dlp OE to mitigate age dependent axonal thinning, restore TBPH localization to the nucleus or improve lifespan and found no significant effects exerted by (G4C2)_44X_ expression in MBNs on these phenotypes (data not shown). These results indicate that Dlp co-OE mitigates some, but not all aspects of *C9orf72*-associated phenotypes and suggest that multiple mechanisms are at play to mediate the toxic effects of G4C2 HR expansions in ALS/FTD.

**Figure 7.**
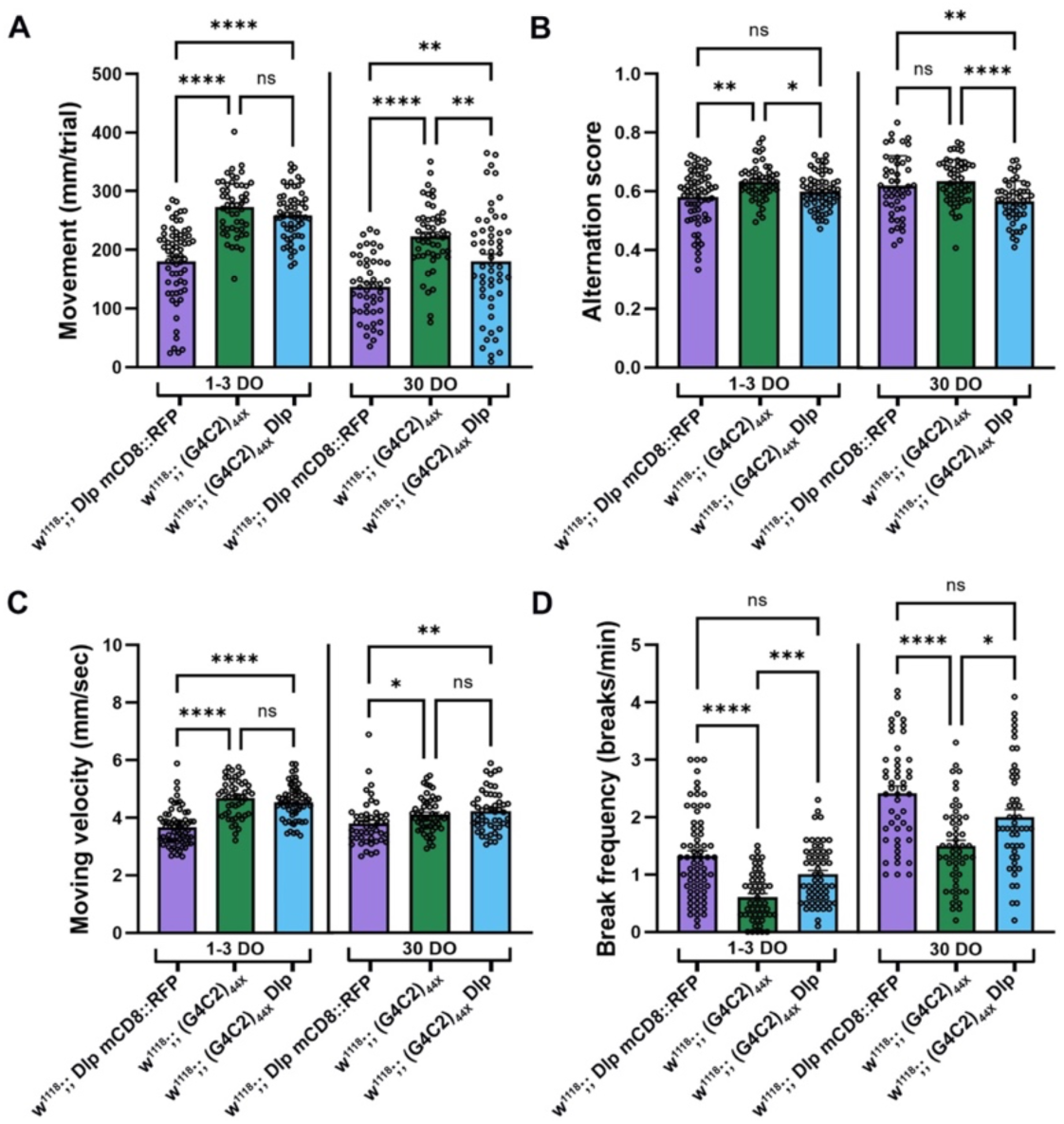
Restoration of Dlp mitigates locomotor deficits and spatial memory alterations in (G4C2)_44X_ *Drosophila* disease model. Y-maze measurements of **(A)** Movement, **(B)** Alternation scores indicating spontaneous alternations. **(C)** Moving velocity, and **(D)** Break frequency. Genotypes and ages, as indicated. N≈35 flies/sex/genotype/timepoint. Significance was calculated using Kruskal-Wallis with Dunn’s multiple-comparison test; *P < 0.05, **P < 0.01, ***P < 0.001, ****P < 0.0001.

### G4C2 expression in *Drosophila* MBs causes a reduction in presynaptic active zones that is mitigated by Dlp overexpression in an age specific manner

Previous studies in flies and zebrafish identified *C9orf72*-associated deficits in synaptic vesicle cycling and presynaptic active zones [70, 71]. To assess whether G4C2 HR expression affects presynaptic active zones (AZs) in MBNs, we examined the distribution of Bruchpilot (BRP), an established AZ marker [46]. Immunofluorescence experiments showed that BRP signal intensity was significantly reduced in γ lobes of (G4C2)_44X_ expressing flies compared to w^1118^ controls at 30 DO (see arrowheads, **Figure 8 A-A3**, **C-C3**, **F**; P = 0.0007). Next, we evaluated the ability of Dlp to mitigate the AZ reduction caused by (G4C2)_44X_ by overexpressing Dlp. These experiments showed that restoring Dlp expression in the context of (G4C2)_44X_ resulted in a partial rescue of presynaptic AZs compared to (G4C2)_44X_ alone, as indicated by BRP intensity in the γ lobe (see arrowheads, **Figure 8 D-D3**, **F;** P = 0.0031). No significant differences were found between presynaptic AZs in Dlp co-OE with (G4C2)_44X_ and w^1118^ (P > 0.9999) or Dlp OE alone (P > 0.9999). Similarly, Dlp OE AZs were comparable with w^1118^ (P = 0.5346) but were significantly higher than (G4C2)_44X_ (P = 0.0403). Interestingly, a significant reduction in AZs was observed in (G4C2)_44X_ HR compared to controls in 1-3 DO brains (**Figure 8E**; P = 0.0329 and data not shown), however Dlp OE did not mitigate this phenotype (**Figure 8E**; P > 0.9999 and data not shown). Taken together these findings support the notion that G4C2 repeat expression in *Drosophila* causes progressive loss of presynaptic active zones in MB axons and restoring Dlp expression can mitigate this phenotype in an age dependent manner.

**Figure 8.**
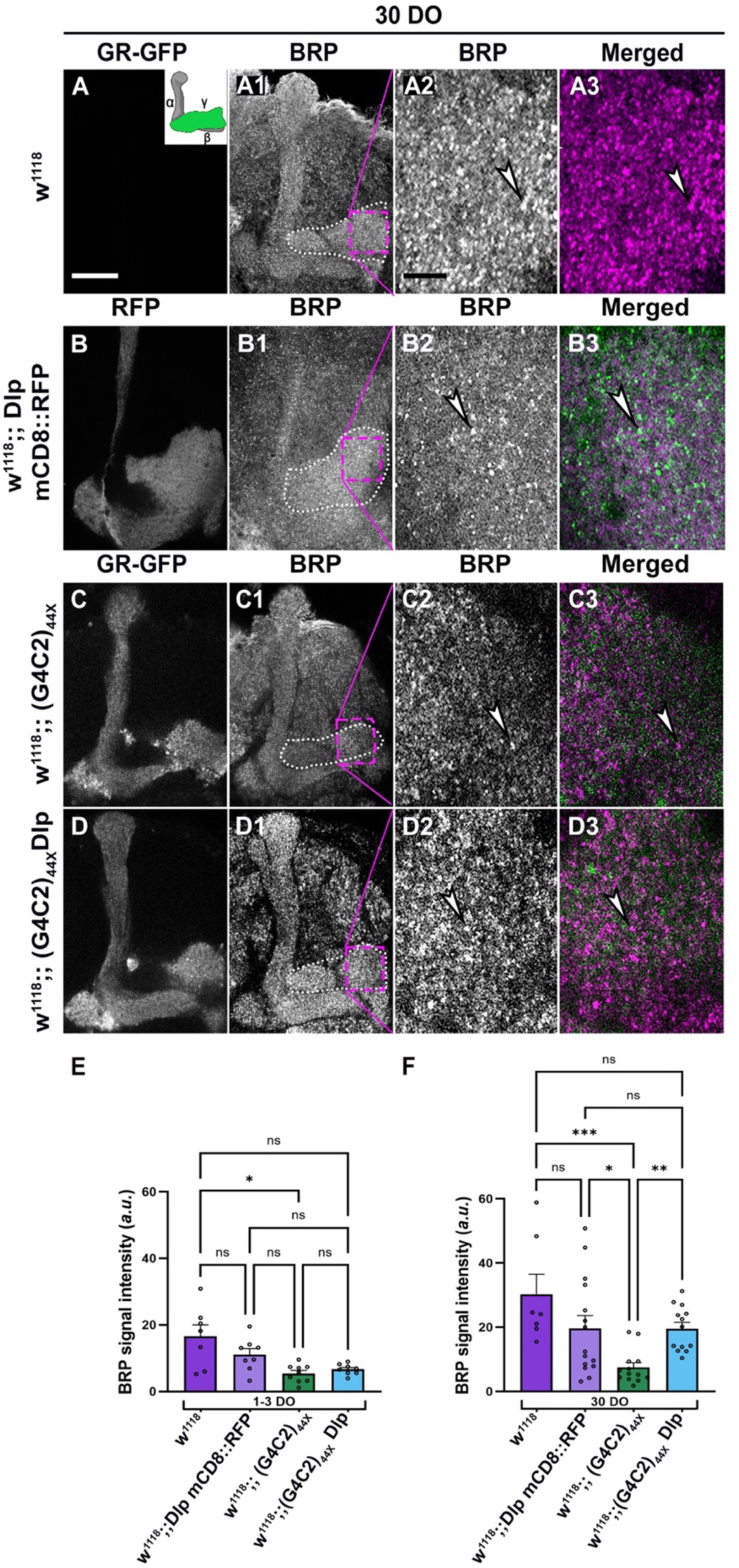
Dally-like protein (Dlp) restores G4C2 HRs-induced loss of presynaptic active zones in MB γ lobe axons. (A-D3) Confocal images showing GR-GFP and BRP expression in MB γ lobe axons (white dotted contour) of (G4C2)_44X_ **(C-C3)**, w^1118^ background control **(A-A3),** Dlp co-OE with RFP (**B-B3**) a transgene control, and Dlp co-OE with (G4C2)_44X_ (**D-D3**) flies at 30 DO **(A-D3)**. **(A2, A3, B2, B3, C2, C3, and D2, D3)** high magnification views of insets shown by magenta lines **(A1-D1)**. **(E, F)** Quantification of BRP signal intensity in the MB γ lobe is shown for 1-3 and 30 DO flies. Scale bars: 50 µm in **A** and 5 µm in **A2**. Genotypes and age, as indicated. N≈7-15 brains/genotype. Data were analyzed using the Kruskal-Wallis with Dunn’s Multiple Comparison Test; *P < 0.05, **P < 0.01, ***P < 0.001.

### GPC6, a human ortholog of Dlp, synergizes with TDP-43 to support synapse formation and maintenance in human iNeurons

Next, we sought to determine whether loss of GPC6, a human ortholog of Dlp, plays a role in synapse formation and maintenance in human neurons, in the context of *C9orf72* FTD associated pathology. We used TDP-43 loss of function as a proxy for TDP-43 proteinopathy associated with *C9orf72* pathology [72], which was also observed in our model. Indeed, reducing TDP-43 levels with TARDBP siRNA caused a significant decrease in neurite length (P = 0.0067) and synaptic density (P = 0.0024) (**Figures 9A, B**). This is consistent with recent studies showing that loss of TDP-43 leads to loss of transcripts encoding genes important for synapse formation, function, or maintenance such as *STMN2*, *UNC13A*, and *SYT1* [73, 74]. Next, we established a rabies virus–based assay to quantify synaptic connectivity in NGN2-induced excitatory neurons (iNs) [75], which readily form synapses in culture (**Figure 9C**). In this system, when rabies virus infects a neuron that is a postsynaptic cell, it can travel retrogradely to the presynaptic cell and express a reporter gene. Using this approach, we found that rabies virus encoding a GFP cassette could label presynaptic neurons, providing a method for quantifying structurally intact synapses in iN cultures (**Figure 9C**). Reducing TDP-43 levels with TARDBP siRNA in this system resulted in a significant decrease in GFP-labeled presynaptic neurons (**Figure 9D**). We next leveraged this assay in a CRISPR interference (CRISPRi) screen designed to identify genes influencing synapse number. Using a dCas9-KRAB system, we suppressed a set of candidate genes in iNs while quantifying presynaptic inputs via the rabies virus–mediated GFP labeling approach. Strikingly, sgRNAs targeting GPC6 emerged as strong modifiers in this screen. In the presence of TARDBP siRNA, GPC6-targeting sgRNAs were significantly depleted at both the individual sgRNA and gene levels, specifically in GFP(+) mCherry(-) presynaptic neurons (**Figures 9E, F**) suggesting that knocking down GPC6 further impairs synapse formation, whereas sgRNAs targeting housekeeping genes were unaffected. These data suggest that loss of GPC6 has a detrimental effect on synapse number when TDP-43 levels are reduced, implying that the *C9orf72* associated reduction in GPC6 synergizes with TDP-43 loss of function mechanisms to cause an even greater reduction of synapses. Thus, two different processes resulting from *C9orf72* associated pathology may ultimately lead to synaptic loss in ALS/FTD.

**Figure 9.**
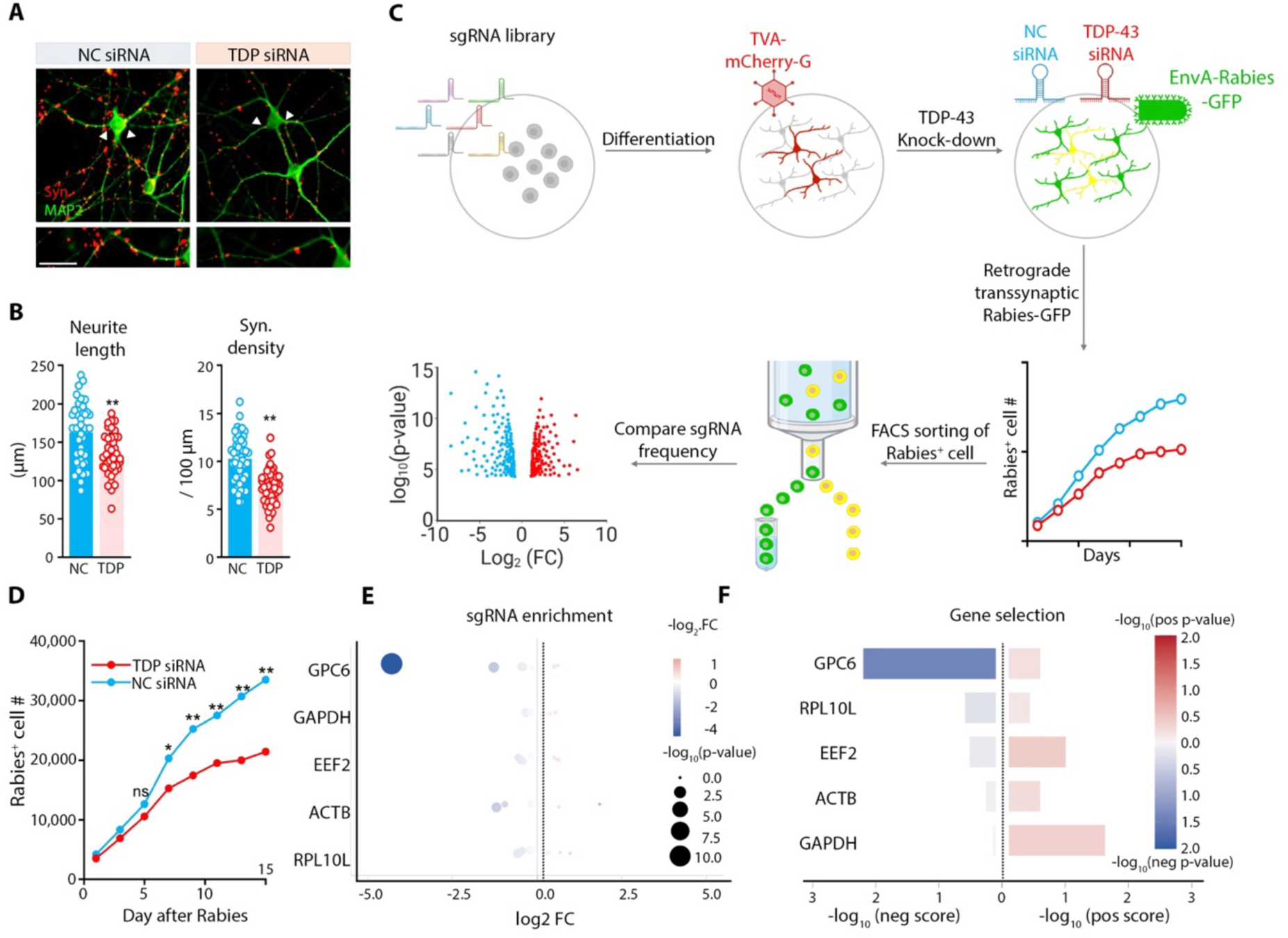
GPC6 siRNA knock-down synergizes with TDP-43 loss of function to regulate synapse numbers in iNeurons. **(A)** Images of Synapsin+ synaptic puncta on the neurites of iNs in cultures treated with a non-targeting (NC) or *TARDBP* (TDP) siRNA. **(B)** Quantification of neurite length and synaptic density on iNs in cultures treated with a non-targeting (NC) or *TARDBP* (TDP) siRNA. **(C)** Overview of rabies virus-labeling of pre-synaptic iNs that form synapses. TVA-mCherry-G are “starter” neurons that can be infected with Rabies-GFP while GFP(+) mCherry(-) iNs are presynaptic neurons that form contacts with the mCherry (+) “starter” cells. **(D)** Number of GFP(+) mCherry(-) pre-synaptic iNs at different days after rabies transduction with treatment with a TARDBP (TDP-43) or non-targeting siRNA. Data shows significant reduction in GFP-labeled neurons at day 7(P = 0.02), day 9 (P = 0.014), day 11(P = 0.0023), day 13 (P = 0.0011) and day 15 (P = 0.0013). **(E)** Each dot represents one sgRNA. The x-axis and dot color show sgRNA-level log2 fold change (LFC) of normalized read counts between TARDBP and non-targeting siRNA. Dot size represents significance of LFC, shown as −log10(p-value). P-value was estimated using MAGeCK’s negative binomial–based count model to test sgRNA depletion/enrichment in treatment relative to control. **(F)** Gene-level selection was analyzed using MAGeCK robust rank aggregation (RRA). Bar plot indicates RRA score for negative or positive enrichment of all sgRNAs targeting each gene. Neg scores (GCP6: 0.008, RPL10L: 0.323, EEF2: 0.383, ACTB: 0.698 and GAPDH: 0.895) suggest sgRNA depletion of a gene while positive scores (GCP6: 0.318, RPL10L: 0.463), EEF2: 0.125, ACTB: 0.318 and GAPDH: 0.03) suggest sgRNA enrichment. Bar color indicates gene-level significance for sgRNA depletion or enrichment. Negative P-value (GCP6: 0.041, RPL10L: 0.602, EEF2: 0.737, ACTB: 0.863 and GAPDH: 0.98) or positive P-value (GCP6: 0.606, RPL10L: 0.725, EEF2: 0.403, ACTB: 0.584 and GAPDH: 0.895) tests whether sgRNAs targeting each gene were randomly enriched among the most depleted (negative) or enriched (positive) sgRNAs. *P < 0.05, **P < 0.01.

## DISCUSSION

Our findings demonstrate that targeted expression of *C9orf72*-associated G4C2 HRs in the *Drosophila* MB circuit recapitulates multiple, age-dependent hallmarks of ALS/FTD pathology. We show that expanded (G4C2)_44X_ repeats drive progressive axonal degeneration, accumulation of RAN-translated GR-GFP dipeptide puncta, and premature nuclear-to-cytoplasmic mislocalization of endogenous TDP-43 (TBPH), accompanied by increased caspase signaling in MBNs.

Within the MB circuit, γ lobe axons show significant thinning in the context of (G4C2)_44X_ HR earlier than α or β lobes, suggesting that these earlier-born MBNs are more significantly impacted and highlighting the aging component of *C9orf72*-associated ALS/FTD. Surprisingly, the number of MBNs is not changed significantly across age (data not shown), yet the axonal lobes are thinner, supporting a “dying back” scenario, which is bolstered by an age-dependent significant reduction in presynaptic active zones [reviewed in 76], and our finding that G4C2 expression leads to a loss of presynaptic active zones in MBNs. This observation deepens the linking of repeat toxicity to synaptic dysfunction [70, 71, 77], and supports proposed models in which repeat mediated toxicity disrupts synaptic function during disease progression, compromises circuit function, and contributes to progressive axonal degeneration [78]. At this time we cannot rule out that G4C2 HRs cause axonal thinning via alterations in memory acquisition and/or consolidation, both of which are known to impact plasticity in MBNs [79].

While the lack of significant changes in overall MBN numbers seemingly contradicts the significant increase in DCP1 signal with age and in a (G4C2)_44X_ dependent manner, it is possible that a caspase cascade is activated by HR expression, however, this may not be sufficient to result in cell death and clearance. Indeed, while DCP1 is considered an “executioner” caspase, some cells require activation of additional caspases for apoptosis [80], which suggests that neuronal cell death involves more complex signaling mechanisms. Notably, γ lobes are thinner in (G4C2)_44X_ before any notable differences in DCP1 signal, suggesting that HRs impact axonal health and integrity before cell bodies, further supporting the “dying back” mechanism of degeneration [reviewed in 76].

We discovered that even the shorter (G4C2)_12X_ repeats exhibited age-dependent RAN translation and intermediate pathological phenotypes, highlighting the vulnerability of the MB circuit to the toxicity of repeat lengths traditionally considered benign in other cell types. Previous work with this shorter repeat in *Drosophila* retina has not shown toxicity [27], consistent with tissue- and/or cell-type-specific vulnerabilities associated with different repeat lengths. The lack of axonal thinning in the 12X compared to the 44X may be caused by differences in pathology that repeat length can confer, which has been shown in other experimental systems and patient tissues [4, 81]. Qualitatively, the character of the 12X vs the 44X GR-GFP puncta seems distinct, with the larger repeat appearing to form larger, more dense puncta. This observation should be tested further, as should the differential solubility of these cytoplasmic accumulations. Given the axonal- and active-zone-driven pathology by the 44X repeat, it would be of interest to explore the effects of the repeats in cell types with and without axonal projections. Regardless of the mechanisms, the difference in axonal thinning between 12X and 44X suggests that the repeats themselves have multiple facets of toxicity, including those that go beyond axonal dying back and dysfunction.

Although our experiments do not directly test the contribution of RNA versus DPRs through the use of specially engineered constructs that only generate RNA or DPRs [82], we note that disease-related phenotypes such as effects on GR-GFP puncta, TBPH mislocalization, lifespan, and sleep, were not observed in the (G4C2)_12X_ flies until after the detection of the RAN translation product. These findings suggest that the toxicity of the DPRs drives disease and are consistent with a recent report demonstrating the critical role of DPRs in *C9orf72*-mediated toxicity [83]. To our knowledge, this is the first report of age-dependent RAN translation *in vivo*, which will be interesting to probe in other experimental systems and human tissues. Interestingly, anti-DPR antibodies have been detected in healthy elderly individuals [84], consistent with RAN translation occurring in aging humans in the absence of lengthy pathogenic repeats.

The behavioral consequences of repeat expression mirror core clinical features of FTD, including locomotor hyperactivity, impaired spatial working memory, and disrupted sleep architecture. Importantly, these effects were modulated by sex and age. For (G4C2)_12X_ flies, male and female flies show different patterns of daytime and nighttime sleep bout numbers at 60 DO, with female flies showing an increase in bout numbers compared to controls, while males are comparable to their controls. Conversely, at the same timepoint, male 44X repeat flies exhibit increased movement compared to their controls, while their 44X female counterparts are comparable to controls in their total movement. Behaviorally, sex differences would be expected, as many clinical studies indicate that male and female FTD patients vary in their symptoms [85, 86].

Consistent with the involvement of the glypican Dlp in TDP-43 related pathology [41, 68], we identify a G4C2 HR- and age-dependent reduction of Dlp in γ lobe axons, and show that restoring Dlp levels partially mitigates locomotor and working memory deficits without rescuing axonal thinning, TDP-43 mislocalization, or lifespan. This is not surprising, given that many pathways are impacted by G4C2 HR expansions and associated TDP-43 mislocalization, the latter of which may cause alterations in the expression of TDP-43’s own RNA targets. It remains to be determined as to what specific targets mediate the effects of G4C2 on these other phenotypes. Interestingly, GPC6, the mammalian ortholog of Dlp, has been shown to restore nuclear TDP-43 when knocked-down in mice lacking the GPC6 processing enzyme GDE2 [68]. Although Dlp manipulations did not affect TDP-43 localization in MBNs, these findings and our previous reports of Dlp/GPC6 alterations in TDP-43 proteinopathies [41, 67] support the idea that disrupted Dlp/GPC6-mediated Wnt signaling may contribute to neuronal vulnerability in *C9orf72* repeat expansion models. The rescue of some but not all G4C2 associated phenotypes suggests that only a subset of neurons and/or neuronal processes benefit from Dlp expression (*e.g.*, neurons that mediate specific locomotor functions). The identity of neurons responsive to Dlp overexpression could be determined via FACS-assisted bulk RNA seq using Dlp and BRP as markers. We note that some rescues (*e.g.*, movement) do not occur in young, but only in older flies suggesting the presence an age-related compensation that is Dlp-dependent for this specific phenotype.

Bolstering the relevance of our findings in the *Drosophila* model, in which we observed axonal Dlp reduction and nuclear loss of TDP-43, we find that in human iNeurons, GPC6 knock-down synergizes with TDP-43 siRNA to enhance TDP-43 associated synaptic loss. Importantly, our findings that Dlp overexpression mitigates presynaptic active zones in *Drosophila* highlights the physiological significance of Dlp/GPC6 in synaptic formation and/or maintenance in the context of *C9orf72*.

Together, the results of our work establish a circuit-specific, aging-sensitive *Drosophila* model of *C9orf72-*associated dementia and underscore the multifactorial mechanisms including RNA toxicity, DPR accumulation, impaired proteoglycan signaling, sex differences, and synaptic vulnerability that converge to drive neurodegeneration in ALS/FTD.

## Supplemental Information (see attached)

## Authorship Contribution Statement

BSC and MAB designed and performed experiments, analyzed data, wrote the first draft, reviewed, and edited the final draft of the manuscript. PCF performed experiments, analyzed data, wrote R scripts, wrote the first draft, reviewed, and edited the final draft of the manuscript. YH designed and performed experiments, analyzed data and wrote, reviewed and edited the final draft. RKG designed experiments, reviewed, and edited the final draft. RS and KVK-J contributed to experimental design, reviewed, and edited the final draft. JKI designed experiments, analyzed data, reviewed and edited the final draft. DCZ designed experiments, analyzed data, wrote the first draft, reviewed, and edited the final draft. All authors reviewed, edited, and approved the manuscript.

## Acknowledgements

Funding was provided by R01NS091299 and R21NS141011 to DCZ, and DoD PR180487 to RS and KJ. We acknowledge the Bloomington Stock Center and Aso Yoshi (HHMI Janelia Farm) for stocks, and Dr. Fen-Biao Gao (Chan Medical School) for anti-TBPH antibodies. The 1D4, 13G8 and nc82 monoclonal antibodies developed by C. Goodman, P. Beachy and E. Buchner, respectively were obtained from the Developmental Studies Hybridoma Bank, created by the NICHD of the NIH and maintained at The University of Iowa, Department of Biology, Iowa City, IA 52242. We thank William Giang in the Advanced Light Microscopy Core Facility for imaging support (RRID:SCR_022526) and acknowledge the CBS department microscopy facility.

