## Supplemental information for "*C9orf72*-associated G4C2 hexanucleotide repeat expression in *Drosophila* mushroom bodies causes age dependent TDP-43 pathology and dementia relevant phenotypes mediated in part by the glypican Dlp/GPC6"

This file contains data and statistical analyses for all experimental genotypes and controls for both males and females, as indicated in the main text and herein.

Affiliations:

[1]Department of Cell and Biological Systems, Penn State College of Medicine, 500 University Drive Crescent Building C4605, Hershey, PA 17033, USA

[2]Eli and Edythe Broad CIRM Center for Regenerative Medicine and Stem Cell Research, University of Southern California, 1425 San Pablo Street, BCC 307, Los Angeles, CA 90033 USA

[3]Department of Biology, Middlebury College, McCardell Bicentennial Hall, Middlebury, VT 05753, USA

[4]The Translational Genomics Research Institute, 445 N. Fifth Street, Phoenix, AZ 85004, USA

[5]Barrow Neurological Institute, Dignity Health St. Joseph's Medical Center, 2910 North Third Avenue, Phoenix, AZ 85013, USA

[#] These authors contributed equally to the manuscript

**G4C2 hexanucleotide repeat (HR) expression in mushroom body neurons (MBNs) causes age-dependent axonal thinning and GR-GFP puncta accumulation in male and female adult brains**

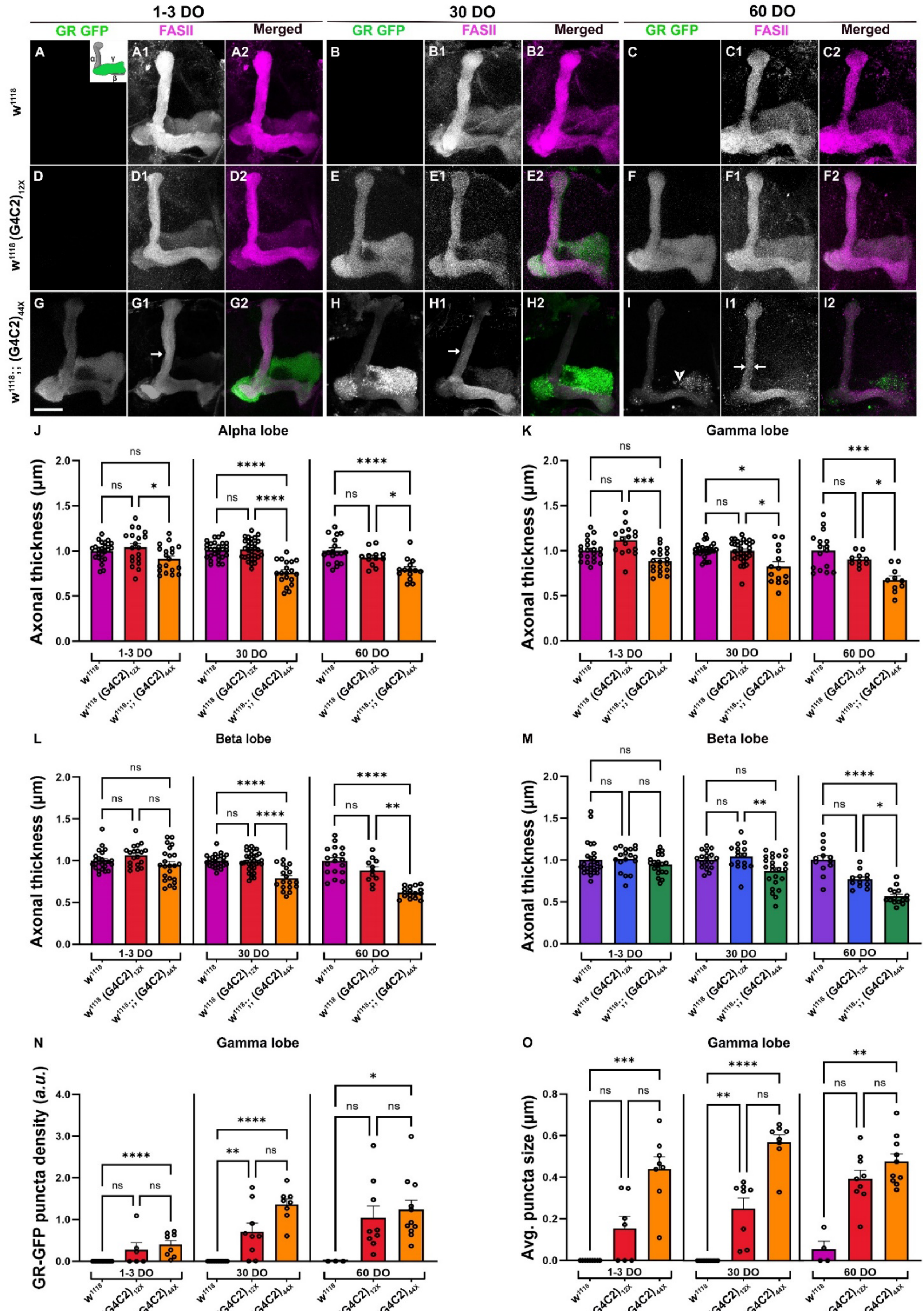

1-3 DO
30 DO
60 DO
GR GFP
FASII
Merged
J
Alpha lobe
K
Gamma lobe
L
Beta lobe
M
Beta lobe
N
Gamma lobe
O
Gamma lobe
Axonal thickness (μm)
GR-GFP puncta density (a.u.)
Avg. puncta size (μm)
ns

**Figure S1. G4C2 HR expression in *Drosophila* MBs causes age-dependent axonal thinning and progressive accumulation of GR-GFP puncta. (A-I2**) $w^{1118}$ controls stained for GR-GFP and FasII. α, β, γ lobe cartoon shown in **(A)**. **(D-F2)** $(G4C2)_{12X}$ stained for GR-GFP and FasII. **(G-I2)** $(G4C2)_{44X}$ stained for GR-GFP and FasII. GR-GFP **(A-I)**, FasII **(A1-I1)** and merged images **(A2-I2)**, as indicated. Arrows indicate thinning α lobes in $(G4C2)_{44X}$ and arrowheads indicate GR-GFP puncta in γ lobes. Scale bar: 50 µm. **(J-L)** Quantification of MB lobes thickness in females, including the α lobe **(J),** γ lobe **(K)** and β lobe **(L),** while **(M)** shows thickness quantification of the β lobe in males. **(N)** Normalized GR-GFP puncta density quantification in 1-3, 30, and 60 DO females. **(O)** Quantification of puncta size in 1-3, 30, and 60 days old (DO) females. Statistical analyses were performed using Kruskal-Wallis with Dunn's multiple-comparison test; $^{ns}P > 0.05$, $*P < 0.05$, $**P < 0.01$, $***P < 0.001$, $****P < 0.0001$.

P-Values:

J) Female Alpha Lobe Thickness

   a. $w^{1118}$ vs $w^{1118}$ $(G4C2)_{12X}$: P >0.9999, 1-3 DO; P >0.9999, 30 DO; P =0.5991, 60 DO
   b. $w^{1118}$ vs $w^{1118}$;; $(G4C2)_{44X}$: P =0.1882, 1-3 DO; P <0.0001, 30 DO; P <0.0001, 60 DO
   c. $w^{1118}$ $(G4C2)_{12X}$ vs $w^{1118}$;; $(G4C2)_{44X}$: P =0.0423, 1-3 DO; P <0.0001, 30 DO; P =0.0326, 60 DO

K) Female Beta Lobe Thickness

   a. $w^{1118}$ vs $w^{1118}$ $(G4C2)_{12X}$: P =0.4401, 1-3 DO; P >0.9999, 30 DO; P =0.6722, 60 DO
   b. $w^{1118}$ vs $w^{1118}$;; $(G4C2)_{44X}$: P >0.9999, 1-3 DO; P <0.0001, 30 DO; P <0.0001, 60 DO
   c. $w^{1118}$ $(G4C2)_{12X}$ vs $w^{1118}$;; $(G4C2)_{44X}$: P =0.0746, 1-3 DO; P <0.0001, 30 DO; P =0.0013, 60 DO

L) Female Gamma Lobe Thickness

   a. $w^{1118}$ vs $w^{1118}$ $(G4C2)_{12X}$: P =0.1131, 1-3 DO; P >0.9999, 30 DO; P >0.9999, 60 DO
   b. $w^{1118}$ vs $w^{1118}$;; $(G4C2)_{44X}$: P =0.0776, 1-3 DO; P =0.0305, 30 DO; P =0.0005, 60 DO
   c. $w^{1118}$ $(G4C2)_{12X}$ vs $w^{1118}$;; $(G4C2)_{44X}$: P =0.0001, 1-3 DO;P =0.0191, 30 DO; P =0.0163, 60 DO

M) Male Beta Lobe Thickness

   a. $w^{1118}$ vs $w^{1118}$ $(G4C2)_{12X}$: P =0.2654, 1-3 DO; P >0.9999, 30 DO; P =0.1039, 60 DO
   b. $w^{1118}$ vs $w^{1118}$;; $(G4C2)_{44X}$: P >0.9999, 1-3 DO; P =0.0998, 30 DO; P <0.0001, 60 DO
   c. $w^{1118}$ $(G4C2)_{12X}$ vs $w^{1118}$;; $(G4C2)_{44X}$: P =0.2956, 1-3 DO; P =0.0072, 30 DO; P =0.0108, 60 DO

N) Female GR-GFP Puncta Density

a. $w^{1118}$ vs $w^{1118}$ $(G4C2)_{12X}$: P =0.0924, 1-3 DO; P =0.0021, 30 DO; P =0.0809, 60 DO

b. $w^{1118}$ vs $w^{1118}$;; $(G4C2)_{44X}$: P <0.0001, 1-3 DO; P <0.0001, 30 DO; P =0.0119, 60 DO

c. $w^{1118}$ $(G4C2)_{12X}$ vs $w^{1118}$;; $(G4C2)_{44X}$: P =0.3363, 1-3 DO; P =0.7095, 30 DO; P >0.9999, 60 DO

O) Female GR-GFP Puncta Size

a. $w^{1118}$ vs $w^{1118}$ $(G4C2)_{12X}$: P =0.3106, 1-3 DO; P =0.0096, 30 DO; P =0.0550, 60 DO

b. $w^{1118}$ vs $w^{1118}$;; $(G4C2)_{44X}$: P =0.0001, 1-3 DO; P <0.0001, 30 DO; P =0.0030, 60 DO

c. $w^{1118}$ $(G4C2)_{12X}$ vs $w^{1118}$;; $(G4C2)_{44X}$: P =0.0671, 1-3 DO; P =0.2621, 30 DO; P =0.7480, 60 DO

**Endogenous dTDP-43 (TBPH) exhibits nuclear to cytoplasmic mislocalization in the context of G4C2 HR expression in Kenyon cells**

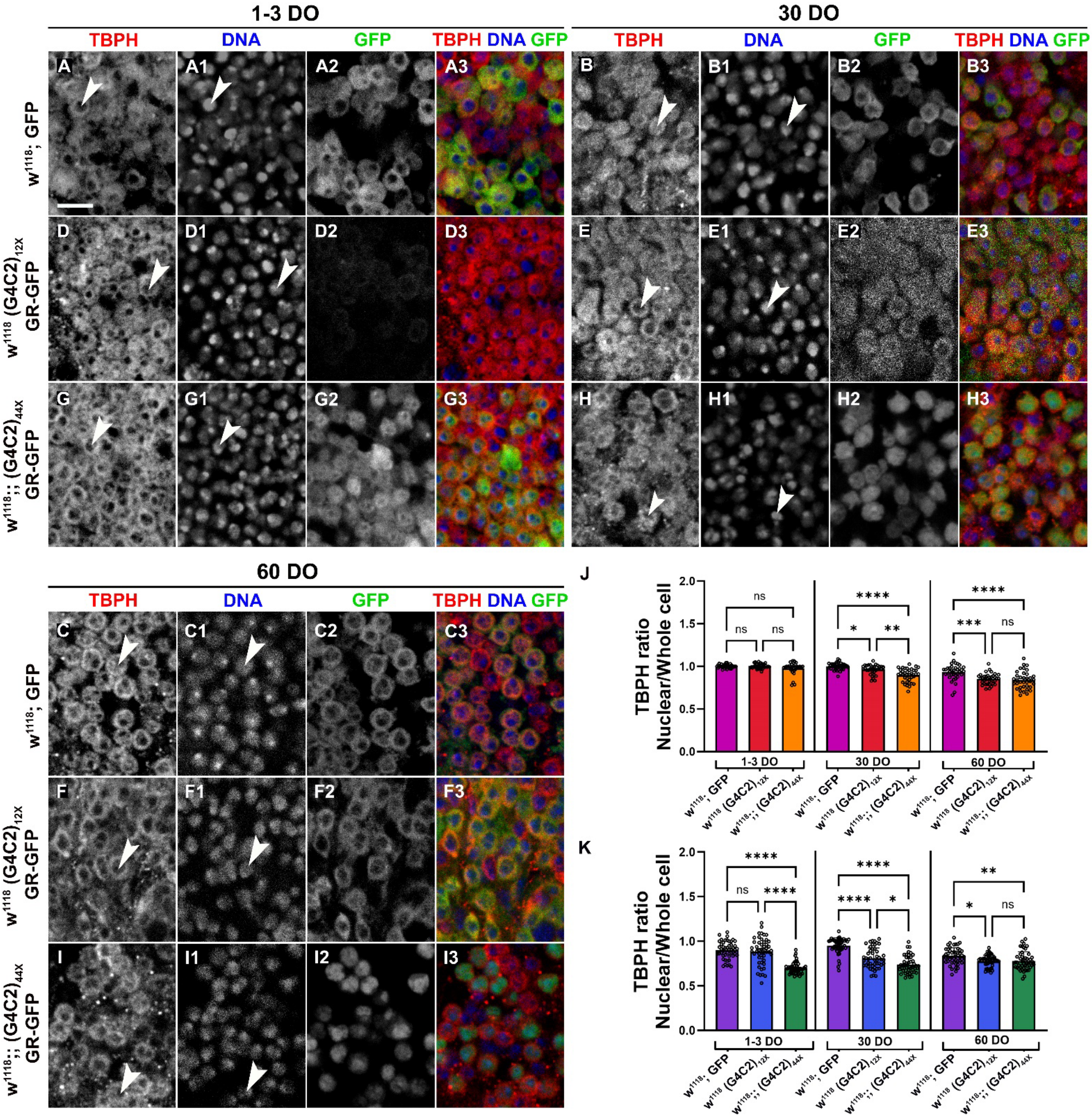


**Figure S2. Expression of G4C2 HR in KCs leads to progressive, age-dependent alterations of endogenous TDP-43 (TBPH) in flies.** Representative confocal images of MBNs are shown at 1-3 DO **(A-A3, D-D3,** and **G-G3)**, 30 DO **(B-B3, E-E3,** and **H-H3)**, and 60 DO **(C-C3, F-F3,** and **I-I3)**. Each age group shows TBPH immunostaining **(A-I)**, nuclear DNA labeling with Hoechst **(A1-I1)**, GFP fluorescence to indicate G4C2 HR expression **(A2-I2)**, and merged images **(A3-I3)**. Scale bar: 5 µm. Quantification of TBPH localization in **(J)** female and **(K)** male KCs across lifespan. Statistical significance was assessed using

Kruskal-Wallis with Dunn's multiple-comparison test; $^{ns}P > 0.05$, $^{*}P < 0.05$, $^{**}P < 0.01$, $^{***}P < 0.001$, $^{****}P < 0.0001$.

P-Values:

J) Female TBPH Nuclear/Whole Cell Ratio

a. $w^{1118}$; GFP vs $w^{1118}$ $(G4C2)_{12X}$: P >0.9999, 1-3 DO; P =0.0247, 30 DO; P =0.0002, 60 DO
b. $w^{1118}$; GFP vs $w^{1118}$;; $(G4C2)_{44X}$: P =0.5054, 1-3 DO; P <0.0001, 30 DO; P <0.0001, 60 DO
c. $w^{1118}$ $(G4C2)_{12X}$ vs $w^{1118}$;; $(G4C2)_{44X}$: P =0.5822, 1-3 DO; P =0.0018, 30 DO; P >0.9999, 60 DO

K) Male TBPH Nuclear/ Whole cell Ratio

a. $w^{1118}$; GFP vs $w^{1118}$ $(G4C2)_{12X}$: P >0.9999, 1-3 DO; P <0.0001, 30 DO; P =0.0112, 60 DO
b. $w^{1118}$; GFP vs $w^{1118}$;; $(G4C2)_{44X}$: P <0.0001, 1-3 DO; P <0.0001, 30 DO; P =0.0036, 60 DO
c. $w^{1118}$ $(G4C2)_{12X}$ vs $w^{1118}$;; $(G4C2)_{44X}$: P <0.0001; 1-3 DO; P =0.0185, 30 DO; P >0.9999, 60 DO

**G4C2 HR expression in MBNs drives age-dependent progressive cell death and reduces lifespan in flies**

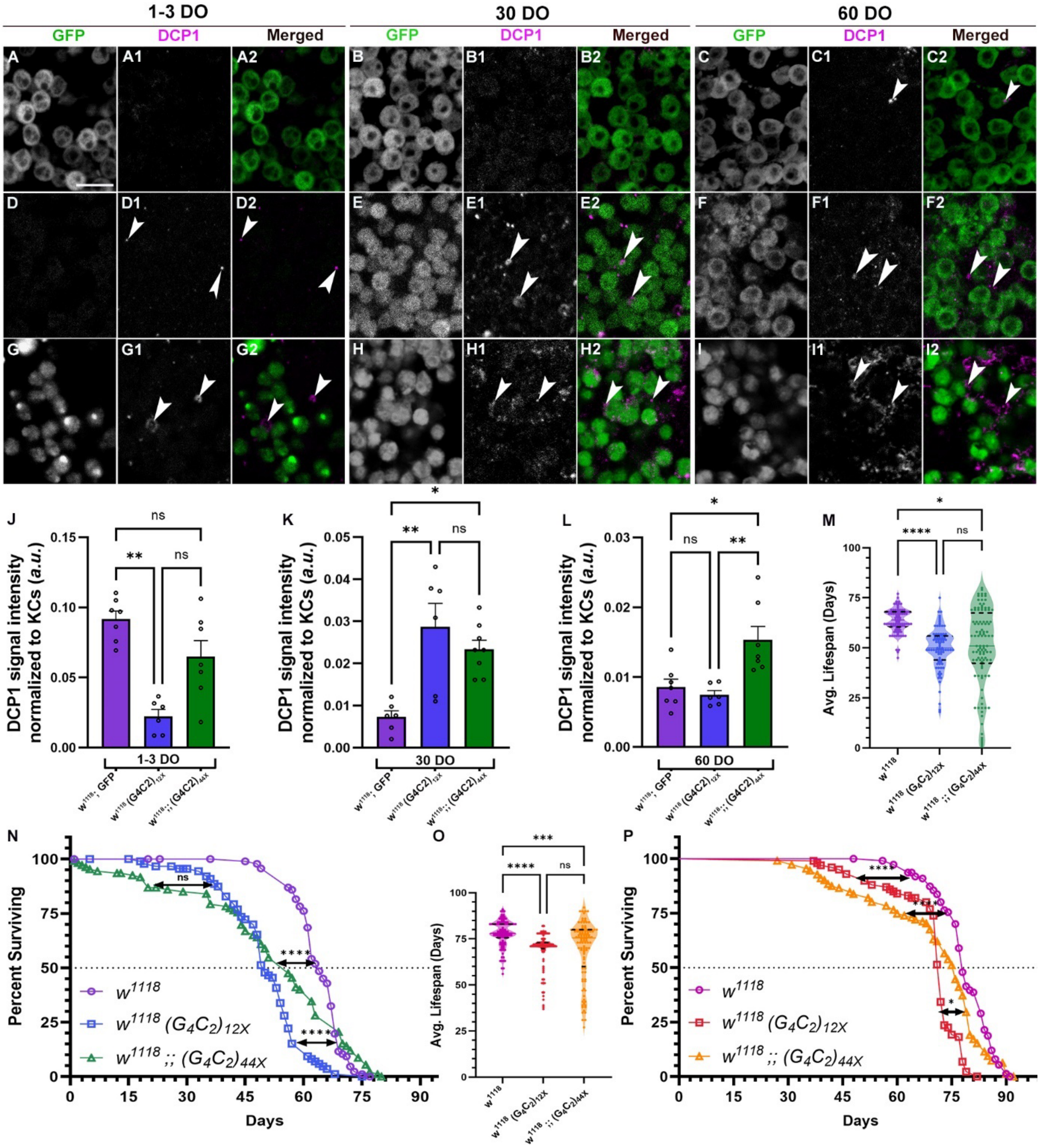


**Figure S3. G4C2 HR expression in *Drosophila* MBNs induces age-dependent apoptotic signaling in Kenyon cells (KCs) and reduces lifespan. (A – I2)** Confocal images showing GFP expression **(A - I)** and cleaved DCP-1 immunostaining **(A1 – I1)** in KCs of GFP, $(G4C2)_{12X}$, and $(G4C2)_{44X}$ expressing female flies. Genotypes, as indicated on the left. Ages, as indicated on the top: 1-3 DO, 30 DO, and 60 DO. Merged images, as shown **(A2 - I2)**. Scale bar: 5 µm. **(J - L)** Quantification of cleaved DCP-1 signal intensity in 1-3

DO **(J)**, 30 DO **(K)**, and 60 DO **(L)** brains. Data were analyzed using Kruskal-Wallis with Dunn's multiple-comparison test; *P < 0.05. **(M – P)** Mean lifespan and survival were analyzed for male and female flies, N ≥ 90 for each genotype/sex. **(M)** Average lifespan for male and **(O)** female flies, for both controls and G4C2 repeat expressing flies in MBNs. Data were analyzed using Kruskal-Wallis with statistical significance assessed using Dunn's test. **(N)** Survival curves for male and **(P)** female flies, for both controls and G4C2 repeat expressing flies in MBNs. The dotted line indicates the 50% survival point (median lifespan). Data were analyzed using the Kaplan-Meier method. Statistical significance was assessed by the Gehan-Breslow-Wilcoxon test; $^{ns}$P >0.05, *P <0.05, **P <0.01, ***P <0.001, ****P <0.0001.

P-Values:

J) Male DCP1 Signal 1-3 DO

a. $w^{1118}$; GFP vs $w^{1118}$ $(G4C2)_{12X}$: P =0.0019
b. $w^{1118}$; GFP vs $w^{1118}$;; $(G4C2)_{44X}$: P =0.4841
c. $w^{1118}$ $(G4C2)_{12X}$ vs $w^{1118}$;; $(G4C2)_{44X}$: P =0.1136

K) Male DCP1 Signal 30 DO

a. $w^{1118}$; GFP vs $w^{1118}$ $(G4C2)_{12X}$: P =0.0045
b. $w^{1118}$; GFP vs $w^{1118}$;; $(G4C2)_{44X}$: P =0.0152
c. $w^{1118}$ $(G4C2)_{12X}$ vs $w^{1118}$;; $(G4C2)_{44X}$: P >0.9999

L) Male DCP1 Signal 60 DO

a. $w^{1118}$; GFP vs $w^{1118}$ $(G4C2)_{12X}$: P >0.9999
b. $w^{1118}$; GFP vs $w^{1118}$;; $(G4C2)_{44X}$: P =0.0264
c. $w^{1118}$ $(G4C2)_{12X}$ vs $w^{1118}$;; $(G4C2)_{44X}$: P =0.0065

M) Male Average Lifespan

a. $w^{1118}$ vs $w^{1118}$ $(G4C2)_{12X}$: P <0.0001
b. $w^{1118}$ vs $w^{1118}$;; $(G4C2)_{44X}$: P =0.0200
c. $w^{1118}$ $(G4C2)_{12X}$ vs $w^{1118}$;; $(G4C2)_{44X}$: P =0.9901

N) Male Survival Curves

a. $w^{1118}$ vs $w^{1118}$ $(G4C2)_{12X}$: P <0.0001

b. $w^{1118}$ vs $w^{1118}$;; $(G4C2)_{44X}$: $P < 0.0001$

c. $w^{1118}$ $(G4C2)_{12X}$ vs $w^{1118}$;; $(G4C2)_{44X}$: $P = 0.0561$

O) Female Average Lifespan

a. $w^{1118}$ vs $w^{1118}$ $(G4C2)_{12X}$: $P < 0.0001$

b. $w^{1118}$ vs $w^{1118}$;; $(G4C2)_{44X}$: $P = 0.0002$

c. $w^{1118}$ $(G4C2)_{12X}$ vs $w^{1118}$;; $(G4C2)_{44X}$: $P > 0.9999$

P) Female Survival Curves

a. $w^{1118}$ vs $w^{1118}$ $(G4C2)_{12X}$: $P < 0.0001$

b. $w^{1118}$ vs $w^{1118}$;; $(G4C2)_{44X}$: $P < 0.0001$

c. $w^{1118}$ $(G4C2)_{12X}$ vs $w^{1118}$;; $(G4C2)_{44X}$: $P = 0.0183$

**G4C2 HR expression in MBNs causes spatial working memory defects in age, sex, and time-dependent manner**

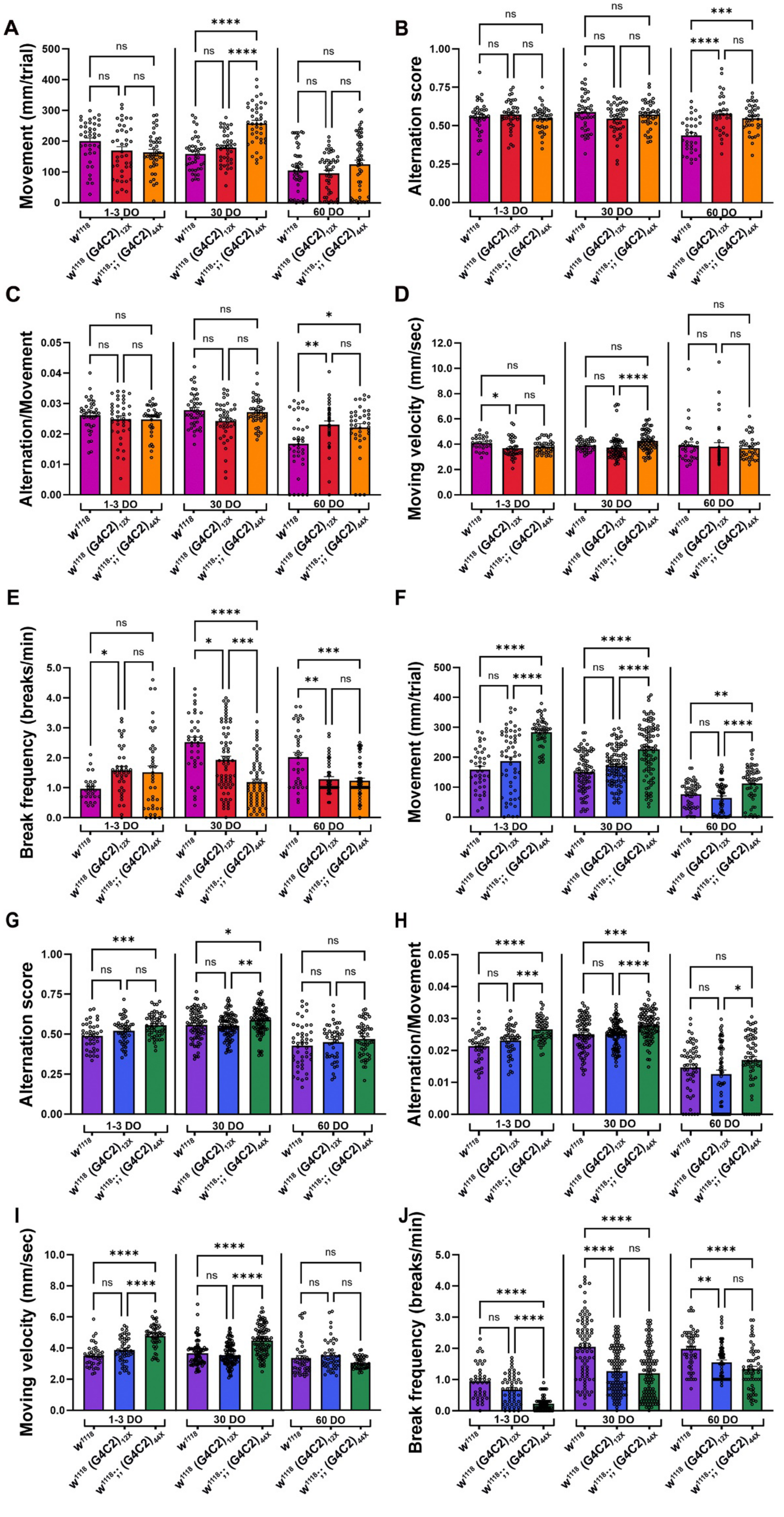

A
Movement (mm/trial)
B
Alternation score
C
Alternation/Movement
D
Moving velocity (mm/sec)
E
Break frequency (breaks/min)
F
Movement (mm/trial)
G
Alternation score
H
Alternation/Movement
I
Moving velocity (mm/sec)
J
Break frequency (breaks/min)
1-3 DO
30 DO
60 DO
w1118
w1118 (G4C2)12X
w1118;; (G4C2)44X
ns
*
**
***
****

**Figure S4. Spatial working memory of $(G4C2)_{44X}$ expressing *Drosophila* at 1-3, 30, and 60 DO.** For female flies, **(A)** Total movement. **(B)** Alternation score indicating spontaneous alternations. **(C)** Alternation numbers normalized to movement. **(D)** Moving velocity, and **(E)** Break frequency. For male flies, **(F)** Total movement. **(G)** Spontaneous alternations presented by the Alternation score. **(H)** Memory performance calculated as number of alternations normalized to movement. **(I)** Average moving velocity, and **(J)** Break frequency during movement. Genotypes and ages are indicated in figure. Each group included N≈35 flies/sex/genotype/timepoint. Data were analyzed using the Kruskal-Wallis with Dunn's multiple-comparison test; $^{ns}P > 0.05$, $^{*}P < 0.05$, $^{**}P < 0.01$, $^{***}P < 0.001$, $^{****}P < 0.0001$.

P-Values:

A) Female Movement

d. $w^{1118}$ vs $w^{1118}$ $(G4C2)_{12X}$: P =0.1763, 1-3 DO; P =0.3605, 30 DO; P >0.9999, 60 DO

a. $w^{1118}$ vs $w^{1118}$;; $(G4C2)_{44X}$: P =0.0545, 1-3 DO; P <0.0001, 30 DO; P =0.6655, 60 DO

b. $w^{1118}$ $(G4C2)_{12X}$ vs $w^{1118}$;; $(G4C2)_{44X}$: P >0.9999, 1-3 DO; P <0.0001, 30 DO; P =0.3408, 60 DO

B) Female Alternation score

a. $w^{1118}$ vs $w^{1118}$ $(G4C2)_{12X}$: P >0.9999, 1-3 DO; P =0.5922, 30 DO; P <0.0001, 60 DO

b. $w^{1118}$ vs $w^{1118}$;; $(G4C2)_{44X}$: P >0.9999, 1-3 DO; P >0.9999, 30 DO; P =0.0003, 60 DO

c. $w^{1118}$ $(G4C2)_{12X}$ vs $w^{1118}$;; $(G4C2)_{44X}$: P =0.6076, 1-3 DO; P >0.9999, 30 DO; P >0.9999, 60 DO

C) Female Alternation/Movement

a. $w^{1118}$ vs $w^{1118}$ $(G4C2)_{12X}$: P >0.9999, 1-3 DO; P =0.0818, 30 DO; P = 0.0068, 60 DO

b. $w^{1118}$ vs $w^{1118}$;; $(G4C2)_{44X}$: P >0.9999, 1-3 DO; P >0.9999, 30 DO; P =0.0129, 60 DO

c. $w^{1118}$ $(G4C2)_{12X}$ vs $w^{1118}$;; $(G4C2)_{44X}$: P =0.1949, 1-3 DO; P =0.1949, 30 DO; P >0.9999, 60 DO

D) Female Moving Velocity

a. $w^{1118}$ vs $w^{1118}$ $(G4C2)_{12X}$: P =0.0108, 1-3 DO; P =0.0623, 30 DO; P =0.6386, 60 DO

b. $w^{1118}$ vs $w^{1118}$;; $(G4C2)_{44X}$: P =0.1985, 1-3 DO; P =0.2618, 30 DO; P >0.9999, 60 DO

c. $w^{1118}$ $(G4C2)_{12X}$ vs $w^{1118}$;; $(G4C2)_{44X}$: P =0.7069, 1-3 DO; P <0.0001, 30 DO; P =0.6590, 60 DO

E) Female Break Frequency

a. $w^{1118}$ vs $w^{1118}$ $(G4C2)_{12X}$: P =0.0191, 1-3 DO; P =0.0419, 30 DO; P =0.0012, 60 DO

b. $w^{1118}$ vs $w^{1118}$ $(G4C2)_{44X}$: P =0.3569, 1-3 DO; P <0.0001, 30 DO; P =0.0004, 60 DO

c. $w^{1118}$ $(G4C2)_{12X}$ vs $w^{1118}$;; $(G4C2)_{44X}$: P =0.5732, 1-3 DO; P =0.0003, 30 DO; P >0.9999, 60 DO

F) Male Movement

a. $w^{1118}$ vs $w^{1118}$ $(G4C2)_{12X}$: P =0.1607, 1-3 DO; P =0.1817, 30 DO; P =0.7413, 60 DO

b. $w^{1118}$ vs $w^{1118}$;; $(G4C2)_{44X}$: P <0.0001, 1-3 DO; P <0.0001, 30 DO; P =0.0020, 60 DO

c. $w^{1118}$ $(G4C2)_{12X}$ vs $w^{1118}$;; $(G4C2)_{44X}$: P <0.0001, 1-3 DO; P <0.0001, 30 DO; P <0.0001, 60 DO.

G) Male Alternation score

a. $w^{1118}$ vs $w^{1118}$ $(G4C2)_{12X}$: P =0.2331, 1-3 DO; P >0.9999, 30 DO; P =0.9999, 60 DO

b. $w^{1118}$ vs $w^{1118}$;; $(G4C2)_{44X}$: P =0.0009, 1-3 DO; P =0.0496, 30 DO; P =0.3176, 60 DO

c. $w^{1118}$ $(G4C2)_{12X}$ vs $w^{1118}$;; $(G4C2)_{44X}$: P =0.1588, 1-3 DO; P =0.0031, 30 DO; P >0.9999, 60 DO

H) Male Alternation/Movement

a. $w^{1118}$ vs $w^{1118}$ $(G4C2)_{12X}$: P =0.2360, 1-3 DO; P >0.9999, 30 DO; P >0.9999, 60 DO

b. $w^{1118}$ vs $w^{1118}$;; $(G4C2)_{44X}$: P <0.0001, 1-3 DO; P =0.0007, 30 DO; P =0.3542, 60 DO

c. $w^{1118}$ $(G4C2)_{12X}$ vs $w^{1118}$;; $(G4C2)_{44X}$: P =0.0009, 1-3 DO; P <0.0001, 30 DO; P =0.0287, 60 DO

I) Male Moving Velocity

a. $w^{1118}$ vs $w^{1118}$ $(G4C2)_{12X}$: P =0.1282, 1-3 DO; P >0.9999, 30 DO; P =0.2831, 60 DO

b. $w^{1118}$ vs $w^{1118}$;; $(G4C2)_{44X}$: P <0.0001, 1-3 DO; P <0.0001, 30 DO; P >0.9999, 60 DO

c. $w^{1118}$ $(G4C2)_{12X}$ vs $w^{1118}$;; $(G4C2)_{44X}$: P <0.0001, 1-3 DO; P <0.0001, 30 DO; P =0.1528, 60 DO

J) Male Break Frequency

a. $w^{1118}$ vs $w^{1118}$ $(G4C2)_{12X}$: P =0.1222, 1-3 DO; P <0.0001, 30 DO; P =0.0026, 60 DO

b. $w^{1118}$ vs $w^{1118}$;; $(G4C2)_{44X}$: P <0.0001, 1-3 DO; P <0.0001, 30 DO; P <0.0001, 60 DO

c. $w^{1118}$ $(G4C2)_{12X}$ vs $w^{1118}$;; $(G4C2)_{44X}$: P <0.0001, 1-3 DO; P >0.9999, 30 DO; P =0.4184, 60 DO

**Expression of G4C2 HR in MBs causes changes in sleep and activity alterations in male and female flies in age-dependent manner**

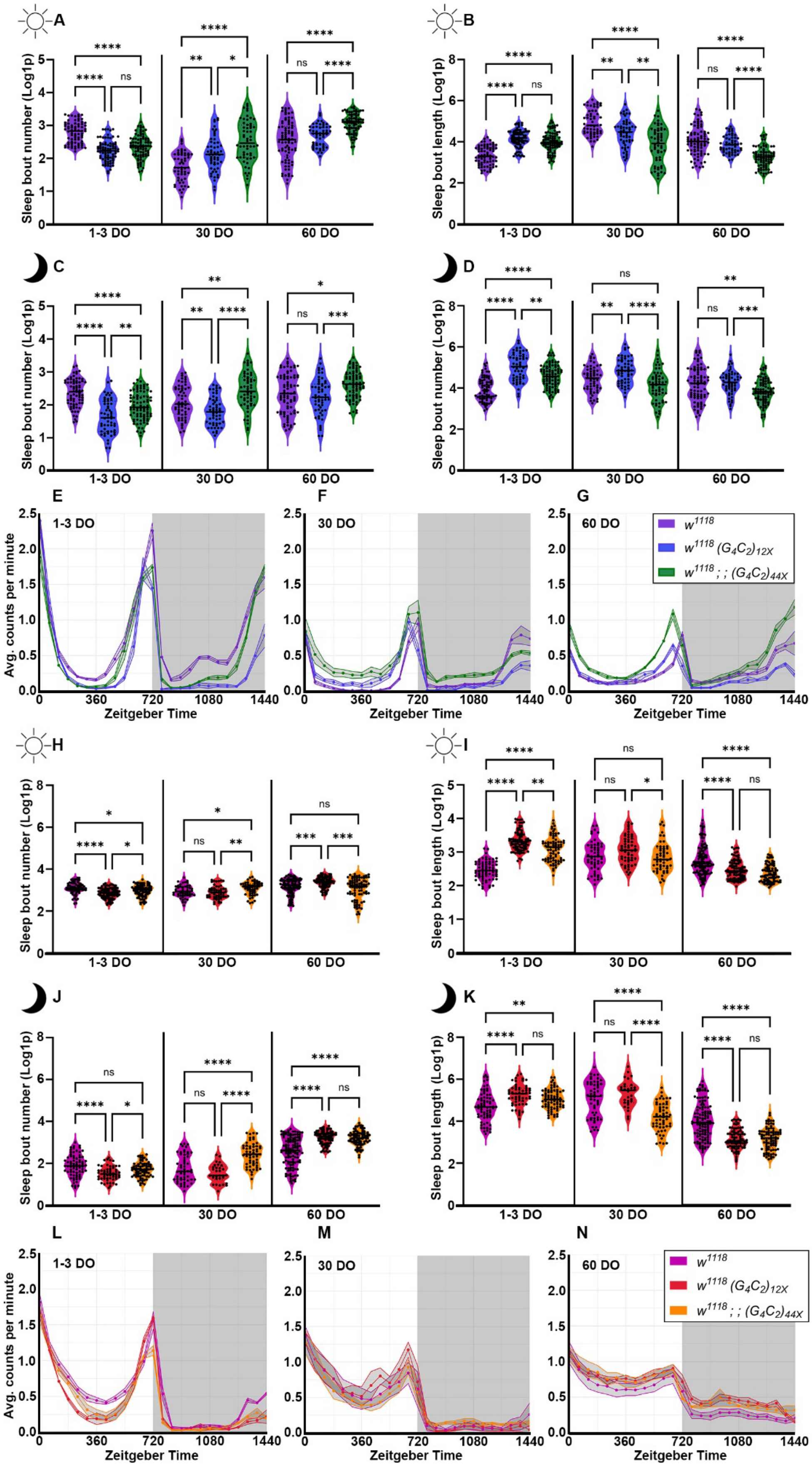

A
B
C
D
E
F
G
H
I
J
K
L
M
N
Sleep bout number (Log1p)
Sleep bout length (Log1p)
1-3 DO
30 DO
60 DO
Avg. counts per minute
Zeitgeber Time
w1118
w1118 (G4C2)12X
w1118 ; ; (G4C2)44X

**Figure S5. Sleep behavior and activity patterns are altered by the expression of G4C2 HR. (A - G)** show male and **(H - N)** female flies, sleep and activity, for control and G4C2-expressing flies, genotypes as indicated in the keys, ages indicated on graphs. Average male daytime sleep bout **(A)** numbers and **(B)** lengths, and male nighttime sleep bout **(C)** numbers and sleep bout **(D)** lengths were recorded during the 12-hour light period (daytime) and 12-hour dark period (nighttime)**. (E-G)** The average number of beam breaks across zeitgeber time for 1-3 DO **(E)**, 30 DO **(F)**, and 60 DO **(G)** male flies. Nighttime indicated by a gray background. Average female daytime sleep bout **(H)** numbers and **(I)** lengths, and male nighttime sleep bout **(J)** numbers and sleep bout **(K)** lengths were recorded during the 12-hour light period (daytime) and 12-hour dark period (nighttime)**. (L - N)** The average number of beam breaks across zeitgeber time for 1-3 DO **(L)**, 30 DO **(M)**, and 60 DO **(N)** male flies. Each group included N≈32 flies/genotype/age/sex. Significance for bouts was determined using a Kruskal-Wallis ANOVA with Dunn's test for multiple comparisons; $^{ns}$P >0.05, *P <0.05, **P <0.01, ***P <0.001, ****P <0.0001.

P-Values:

A) Male Sleep Bout Number Light
   a. $w^{1118}$ vs $w^{1118}$ $(G4C2)_{12X}$: P <0.0001, 1-3 DO; P =0.0014, 30 DO; P >0.9999, 60 DO
   b. $w^{1118}$ vs $w^{1118}$;; $(G4C2)_{44X}$: P <0.0001, 1-3 DO; P <0.0001, 30 DO; P <0.0001, 60 DO
   c. $w^{1118}$ $(G4C2)_{12X}$ vs $w^{1118}$;; $(G4C2)_{44X}$: P =0.0778, 1-3 DO; P =0.0101, 30 DO; P <0.0001, 60 DO
B) Male Sleep Bout Length Light
   a. $w^{1118}$ vs $w^{1118}$ $(G4C2)_{12X}$: P <0.0001, 1-3 DO; P =0.0031, 30 DO; P >0.9999, 60 DO
   b. $w^{1118}$ vs $w^{1118}$;; $(G4C2)_{44X}$: P <0.0001, 1-3 DO; P <0.0001, 30 DO; P <0.0001, 60 DO
   c. $w^{1118}$ $(G4C2)_{12X}$ vs $w^{1118}$;; $(G4C2)_{44X}$: P =0.0772, 1-3 DO; P =0.0061, 30 DO; P <0.0001, 60 DO
C) Male Sleep Bout Number Dark
   a. $w^{1118}$ vs $w^{1118}$ $(G4C2)_{12X}$: P <0.0001, 1-3 DO; P =0.0083, 30 DO; P =0.7800, 60 DO
   b. $w^{1118}$ vs $w^{1118}$;; $(G4C2)_{44X}$: P <0.0001, 1-3 DO; P =0.0099, 30 DO; P =0.0102, 60 DO
   c. $w^{1118}$ $(G4C2)_{12X}$ vs $w^{1118}$;; $(G4C2)_{44X}$: P =0.0069, 1-3 DO; P <0.0001, 30 DO; P =0.0005, 60 DO
D) Male Sleep Bout Length Dark
   a. $w^{1118}$ vs $w^{1118}$ $(G4C2)_{12X}$: P <0.0001, 1-3 DO; P =0.0093, 30 DO; P >0.9999, 60 DO

b. $w^{1118}$ vs $w^{1118}$;; $(G4C2)_{44X}$: P <0.0001, 1-3 DO; P =0.0950, 30 DO; P =0.0071, 60 DO

c. $w^{1118}$ $(G4C2)_{12X}$ vs $w^{1118}$;; $(G4C2)_{44X}$: P =0.0024, 1-3 DO; P <0.0001, 30 DO; P =0.0008, 60 DO

H) Female Sleep Bout Number Light

a. $w^{1118}$ vs $w^{1118}$ $(G4C2)_{12X}$: P <0.0001, 1-3 DO; P >0.9999, 30 DO; P =0.0005, 60 DO

b. $w^{1118}$ vs $w^{1118}$;; $(G4C2)_{44X}$: P =0.0423, 1-3 DO; P =0.0267, 30 DO; P >0.9999, 60 DO

c. $w^{1118}$ $(G4C2)_{12X}$ vs $w^{1118}$;; $(G4C2)_{44X}$: P =0.0186, 1-3 DO; P =0.0015, 30 DO; P =0.0001, 60 DO

I) Female Sleep Bout Length Light

a. $w^{1118}$ vs $w^{1118}$ $(G4C2)_{12X}$: P <0.0001, 1-3 DO; P =0.0707, 30 DO; P <0.0001, 60 DO

b. $w^{1118}$ vs $w^{1118}$;; $(G4C2)_{44X}$: P <0.0001, 1-3 DO; P >0.9999, 30 DO; P <0.0001, 60 DO

c. $w^{1118}$ $(G4C2)_{12X}$ vs $w^{1118}$;; $(G4C2)_{44X}$: P =0.0024, 1-3 DO; P =0.0124, 30 DO; P =0.0577, 60 DO

J) Female Sleep Bout Number Dark

a. $w^{1118}$ vs $w^{1118}$ $(G4C2)_{12X}$: P <0.0001, 1-3 DO; P =0.3485, 30 DO; P <0.0001, 60 DO

b. $w^{1118}$ vs $w^{1118}$;; $(G4C2)_{44X}$: P =0.0594, 1-3 DO; P <0.0001, 30 DO; P <0.0001, 60 DO

c. $w^{1118}$ $(G4C2)_{12X}$ vs $w^{1118}$;; $(G4C2)_{44X}$: P =0.0467, 1-3 DO; P <0.0001, P >0.9999, 60 DO

K) Female Sleep Bout Length Dark

a. $w^{1118}$ vs $w^{1118}$ $(G4C2)_{12X}$: P <0.0001, 1-3 DO; P =0.5386, 30 DO; P <0.0001, 60 DO

b. $w^{1118}$ vs $w^{1118}$;; $(G4C2)_{44X}$: P =0.0049, 1-3 DO; P <0.0001, 30 DO; P <0.0001, 60 DO

c. $w^{1118}$ $(G4C2)_{12X}$ vs $w^{1118}$;; $(G4C2)_{44X}$: P =0.1045, 1-3 DO; P <0.0001, 30 DO; P >0.9999, 60 DO

### G4C2 HR expression does not alter the Dlp expression in Kenyon cells (KCs)

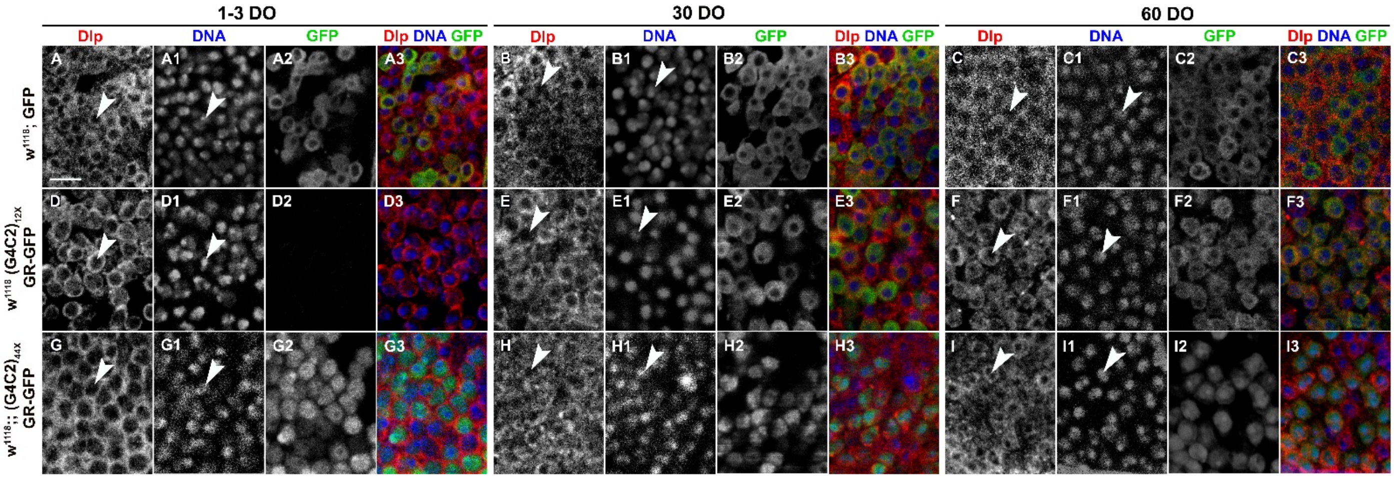


**Figure S6a. Age-dependent Dlp expression in KCs is unaffected by G4C2 HR expansion**. Representative confocal images showing Dlp expression in KCs of male flies at 1-3 DO **(A, D, G)**, 30 DO **(B, E, H)**, and 60 DO **(C, F, I)**. Nuclear DNA labeling with Hoechst at 1-3 DO **(A1, D1, G1)**, 30 DO **(B1, E1, H1)**, and 60 DO **(C1, F1, I1)**. GFP fluorescence to indicate G4C2 HR expression at 1-3 DO **(A2, D2, G2)**, 30 DO **(B2, E2, H2)**, and 60 DO **(C2, F2, I2)**, and merged images at 1-3 DO **(A3, D3, G3)**, 30 DO **(B3, E3, H3)**, and 60 DO **(C3, F3, I3)**. Scale bar: 5 µm.

### G4C2 HR expression in MBNs reduces Dlp expression in γ-lobe axons in an age dependent manner

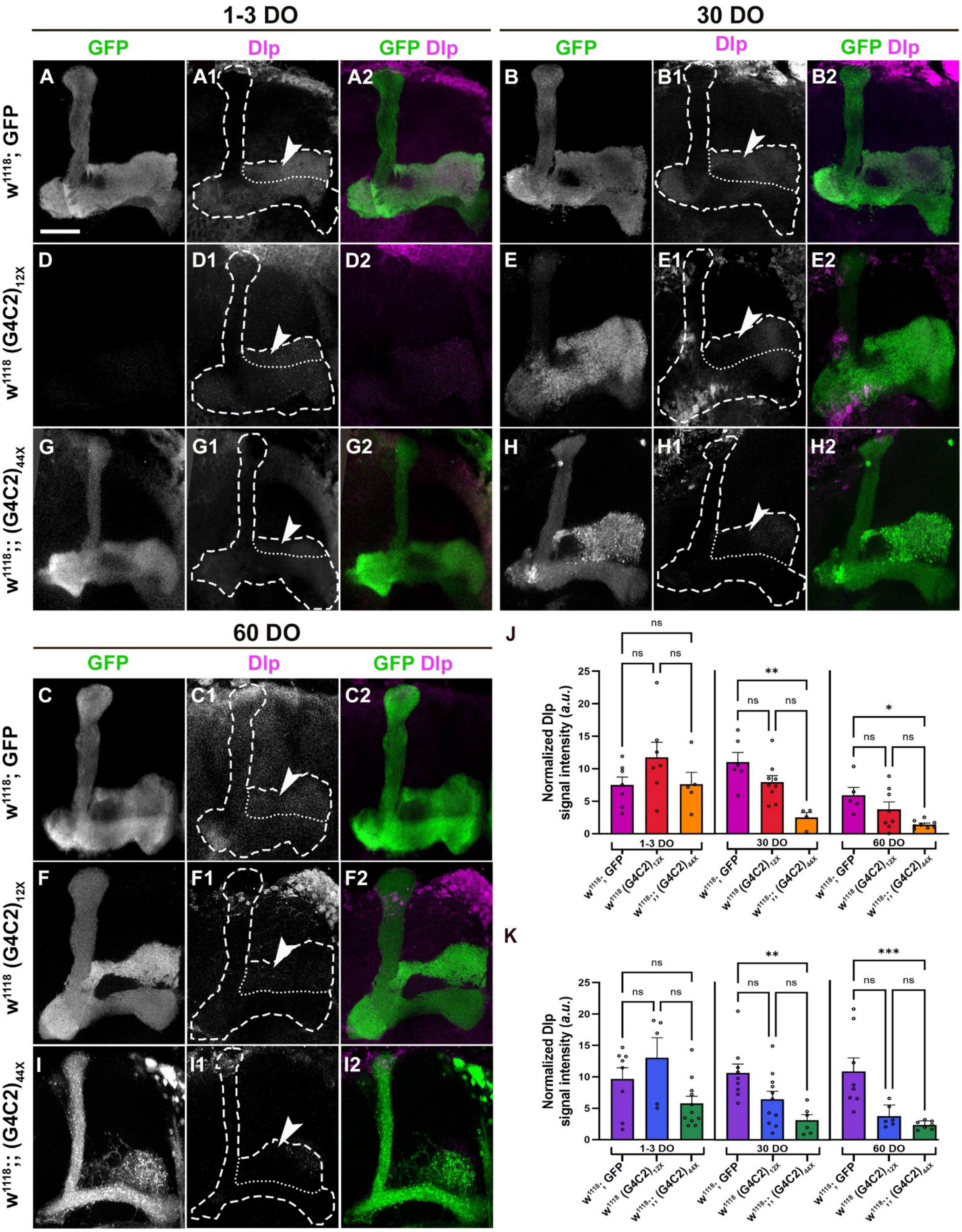

1-3 DO
30 DO
60 DO
GFP
Dlp
GFP Dlp
w1118; GFP
w1118 (G4C2)12X
w1118;; (G4C2)44X
A A1 A2 B B1 B2
D D1 D2 E E1 E2
G G1 G2 H H1 H2
C C1 C2
F F1 F2
I I1 I2
J
K
Normalized Dlp signal intensity (a.u.)
ns
**
*
***

**Figure S6b. Expression of Daly-like protein (Dlp) in the $\gamma$ lobe of MBN is altered in the context of disease G4C2 repeats.** GFP fluorescence for control, and G4C2 HR expressing flies at 1-3 DO **(A, D, G)**, 30 DO **(B, E, H)**, 60 DO **(C, F, I)**. Corresponding Dlp staining is shown at 1-3-days **(A1, D1, G1)**, 30 DO **(B1, E1, H1)**, and 60 DO **(C1, F1, I1)**, with merged images at 1-3 DO **(A2, D2, G2)**, 30 DO **(B2, E2, H2)**, and 60 DO **(C2, F2, I2)**. Scale bar: 50 µm. Quantification of Dlp expression in **(J)** female and **(K)** male $\gamma$ lobe of MB across different age points. Statistical significance was determined using Kruskal-Wallis with Dunn's multiple-comparison test; $^{ns}P > 0.05$, $^{*}P < 0.05$, $^{**}P < 0.01$, $^{***}P < 0.001$.

P-Values:

J) Female Dlp Signal Intensity

a. $w^{1118}$; GFP vs $w^{1118}$ $(G4C2)_{12X}$: P =0.2619, 1-3 DO; P =0.5694, 30 DO; P =0.5449, 60 DO
b. $w^{1118}$; GFP vs $w^{1118}$;; $(G4C2)_{44X}$: P >0.9999, 1-3 DO; P =0.0034, 30 DO; P =0.0144, 60 DO
c. $w^{1118}$ $(G4C2)_{12X}$ vs $w^{1118}$;; $(G4C2)_{44X}$: P =0.3260, 1-3 DO; P =0.0564, 30 DO; P =0.2718, 60 DO

K) Male Dlp Signal Intensity

a. $w^{1118}$; GFP vs $w^{1118}$ $(G4C2)_{12X}$: P =0.9010, 1-3 DO; P =0.1640, 30 DO; P =0.0602, 60 DO
b. $w^{1118}$; GFP vs $w^{1118}$;; $(G4C2)_{44X}$: P =0.5559, 1-3 DO; P =0.0033, 30 DO; P =0.0008, 60 DO
c. $w^{1118}$ $(G4C2)_{12X}$ vs $w^{1118}$;; $(G4C2)_{44X}$: P =0.0761, 1-3 DO; P =0.2730, 30 DO; P =0.7740, 60 DO

**Dlp overexpression does not alter GR-GFP levels in $(G4C2)_{44X}$ flies**

To alleviate concerns about possible dilution effects of GAL4 in the presence of multiple UAS promoters, we performed western blots to evaluate GR-GFP expression in flies expressing $(G4C2)_{44X}$ alone versus co-overexpressing $(G4C2)_{44X}$ and Dlp (Supplemental **Figure S7**).

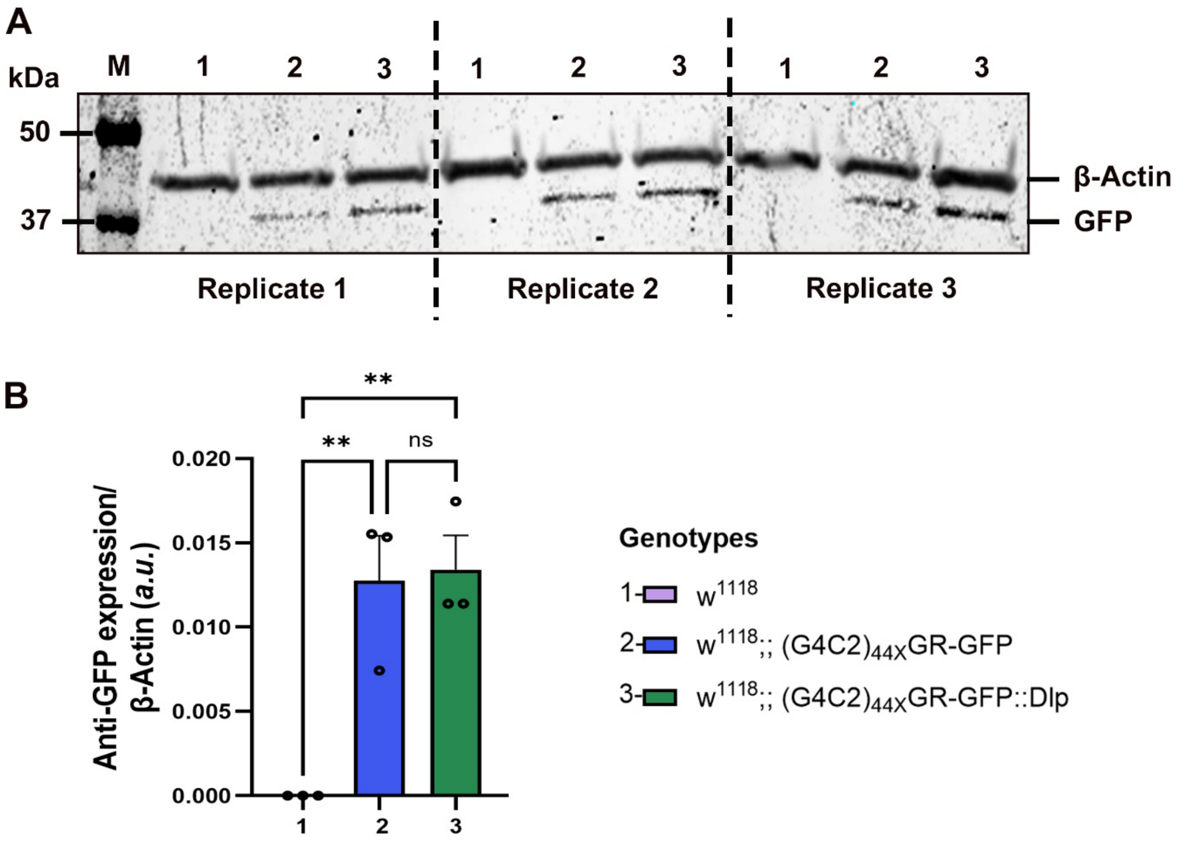


**Figure S7. The expression of GR-GFP is unchanged in the presence of a second UAS-driven transgene. (A)** Western blot exhibits the expression of $w^{1118}$;; $(G4C2)_{44X}$GR-GFP in MBNs, driven by a mushroom body-specific driver line *SS01276.* Blot shown in three different genotypes: $w^{1118}$ background control, $w^{1118}$;; $(G4C2)_{44X}$GR-GFP, and $w^{1118}$;; $(G4C2)_{44X}$GR-GFP::Dlp, with all samples probed using an anti-GFP antibody in the brain of 1-3 DO male flies. The arrows indicate the positions of GFP and β-actin expressions on the right side of the blot. **(B)** The quantification of GFP expression is normalized to β-actin, as shown in **(A)**. The genotypes and replicates are clearly indicated. Statistical significance was determined using ordinary one-way ANOVA test; $^{ns}P > 0.05$, $^{**}P = 0.01$.

P-Values:

A) GFP Expression Using spGAL4

a. $w^{1118}$ vs $w^{1118}$;; $(G4C2)_{44X}$ GR-GFP: P =0.0082, 1-3 DO

b. $w^{1118}$ vs $w^{1118}$;; $(G4C2)_{44X}$GR-GFP:: Dlp: P =0.0065 1-3 DO

c. $w^{1118}$;; $(G4C2)_{44X}$GR-GFP vs $w^{1118}$;; $(G4C2)_{44X}$GR-GFP::Dlp: P =0.9694, 1-3 DO

**Table S1: *Drosophila* lines**

| **Name** | **Collection** | **Identifier** | **Full genotype** | **Ref.** |
|---|---|---|---|---|
| *D. melanogaster:* *-p65ADZp in attP40/CyO;ZpGdbd in attP2* | Janelia FlyLight | SS01276 | *split GAL4* | [1] |
| *D. melanogaster:* $w^{1118}$ | BDSC | RRID: BDSC_5905 | $w^{1118}$ | [2] |
| *D. melanogaster:* $w^{1118}$ *(G4C2)*$_{12X}$ | BDSC | BL# 84722 RRID: BDSC_84722 | *P{w[+mC]=UAS-LDS-(G4C2)12.GR-GFP}2, w[1118]* | [3] |
| *D. melanogaster* $w^{1118}$*;;* *(G4C2)*$_{44X}$ | BDSC | BL# 84723 RRID: BDSC_84723 | *w[1118]; P{w[+mC]=UAS-LDS-(G4C2)44.GR-GFP}9* | [3] |
| *D. melanogaster* $w^{1118}$*;;* *(G4C2)*$_{44X}$ $w^{1118}$*;; dlp* | n/a | n/a | *w[1118]; P{w[+mC]=UAS LDS-(G4C2)44.GR-GFP}9* *w[1118];P{w[+mC]=UAS-dlp.WT}3* | This paper |

| | | | | |
|---|---|---|---|---|
| *D. melanogaster* *mCD8:: RFP* | BDSC | BL# 27399 RRID: BDSC_27399 | *y[1] w[*]; P{w[+mC]=UAS-mCD8.mRFP.LG}10b* | n/a |
| *D. melanogaster* *$w^{1118}$;; dlp* | BDSC | BL# 9160 RRID: BDSC_9160 | *w[1118];P{w[+mC]=UAS-dlp.WT}3* | n/a |
| *D. melanogaster* *$w^{1118}$;; dlp mCD8::RFP* | n/a | n/a | *w[1118]; P{w[+mC]=UAS-dlp.WT}3 y[1] w[*]; P{w[+mC]=UAS-mCD8.mRFP.LG}10b* | This paper |
| *D. melanogaster* *Super GFP* | Robert Kraft | n/a | *w[1118]; UAS super GFP II* | n/a |
| *D. melanogaster* *$w^{1118}$;; $dfmr1^3$* | Tom Jongens | n/a | *w[1118]; P{w[+mC]=UAS dfmr13/TMC6 Sb Tb* | n/a |

**Table S2: List of antibodies/reagents used in these studies.**

| Antibodies/Reagents | Identifier | Dilution | Source/Supplier |
|---|---|---|---|
| Anti-Fasciclin-II | Cat# 1D4 | IF 1:40 | DSHB |

| | | | |
|---|---|---|---|
| | | | RRID: AB_528235 |
| Anti-Dally Like Protein | Cat# 13G8 | IF 1:5 | DSHB<br>RRID: AB_528191 |
| Anti-TBPH | n/a | IF 1:100 | Custom-made in Fen Biao Gao lab, UMass Chan Medical School [4] |
| Anti-Cleaved *Drosophila* Dcp1 (Asp216) | Cat# 9578 | IF 1:500 | Cell Signaling Technology<br>RRID:AB_2721060 |
| Anti-BRP | Cat# NC82 | IF 1:100 | DSHB<br>RRID:AB_2314866 |
| Anti-Synapsin | Cat# 106011 | IF 1:1000 | Synaptic Systems<br>RRID:AB_2619772 |
| anti-MAP2 | Cat# ab5392 | IF: 1: 2000 | Abcam<br>RRID:AB_2138153 |
| Anti-GFP-FITC | Cat# 600-402-215 | IF 1:300 | Rockland<br>RRID:AB_828169 |

| | | | |
|---|---|---|---|
| ChromoTek RFP-Booster ATTO 594 | Cat# rba594 | IF 1:200 | ChromoTek RRID:AB_2631390 |
| Alexa fluor® 488 goat anti-mouse IgG (H+L) | Cat# A11001 | IF 1:500 | Invitrogen RRID:AB_2534069 |
| Alexa fluor® 594 goat anti-rabbit IgG (H+L) | Cat# A11012 | IF 1:500 | Invitrogen RRID:AB_141359 |
| Alexa fluor® 594 goat anti-mouse IgG (H+L) | Cat# A11005 | IF 1:500 | Invitrogen RRID:AB_141372 |
| Alexa fluor® 633 goat anti mouse IgG (H+L) | Cat# A21050 | IF 1:500 | Invitrogen RRID:AB_141431 |
| Alexa fluor® 647 goat anti-rabbit IgG (H+L) | Cat#A21244 | IF 1:500 | Invitrogen RRID:AB_2535812 |
| Hoechst | Cat# 33342 | IF 1:10,000 | Invitrogen RRID:AB_10626776 |
| Anti-GFP (4B10) | Cat# 2955S | WB 1:1000 | Cell Signaling Technology RRID:AB_1196614 |
| Anti-GFP Living Colors Antibody JL-8 | Cat# 632381 | WB 1:1000 | Clontech, A Takara Bio Company RRID:AB_2313808 |

| | | | |
|---|---|---|---|
| β-actin (13E5) | Cat# 93473 | WB 1:1000 | Cell Signaling Technology RRID:AB_3099713 |
